# Machine Learning–Driven Antigen Selection Reveals Conserved T-Cell Targets for Broad Coronavirus Vaccination

**DOI:** 10.64898/2026.04.02.716054

**Authors:** Lorenzo Federico, Alexandru Odainic, Katrine Persgård Lund, Ingrid Marie Egner, Katrin E Wiese, Lisette A H M Cornelissen, Hassen Kared, Richard Stratford, Sebastian Kapell, Brandon Malone, Marius Gheorghe, Pierre Machart, Raman Siarheyeu, Yuki Tanaka, Trevor Clancy, Kaidre Bendjama, Ludvig A. Munthe

## Abstract

**Background:** Coronavirus outbreaks remain a persistent threat to global health, and vaccines based primarily on spike-specific immune responses are susceptible to antigenic variation. T-cell immunity directed against conserved internal viral proteins may provide a complementary and more variant-tolerant strategy for next-generation coronavirus vaccines.

**Methods:** We combined machine learning–guided antigen prioritization with ex vivo functional immunological validation to identify conserved non-spike T-cell targets across betacoronaviruses. Candidate sequences were screened for immunogenicity using primary human peripheral blood mononuclear cells from healthy donors using intracellular cytokine staining and activation-induced marker assays. Top-ranked conserved regions were incorporated into multiepitope mRNA constructs, and their intracellular expression and HLA class I presentation were confirmed by immunopeptidomics. Immunogenicity was further evaluated *ex vivo* and *in vivo* using mRNA immunization of mice and T-cell FluoroSpot assays.

**Findings:** Across a panel of 97 peptides derived from 19 viral proteins, evolutionary conservation across distinct betacoronavirus taxa was strongly associated with functional T-cell immunogenicity in human donors. Highly conserved peptides elicited significantly stronger and more frequent CD4⁺ and CD8⁺ T-cell responses than taxon-restricted peptides. Multiepitope mRNA constructs encoding conserved regions were efficiently expressed and presented on HLA class I molecules and induced T-cell responses in human PBMCs. In mice, mRNA immunization with conserved multiepitope constructs generated robust interferon-γ– and interleukin-2–producing T-cell responses that exceeded those induced by unconserved control constructs.

**Interpretation:** These results link evolutionary conservation to functional cellular immunogenicity and demonstrate the feasibility of multiepitope mRNA delivery for inducing conserved coronavirus-directed T-cell responses. Although protective efficacy remains to be established, conservation-guided antigen selection represents a scalable strategy for developing T-cell–focused vaccines with broad lineage coverage, supporting pandemic preparedness beyond spike-centered immunity.

**Funding:** The research was supported by CEPI, NEC, University of Oslo and Oslo university hospital.

**Research in context:** *Evidence before this study:* Prior coronavirus vaccine development has focused predominantly on spike protein–directed neutralizing antibodies. While highly effective against matched strains, spike-centered immunity is vulnerable to antigenic drift and lineage-specific escape. Multiple observational and experimental studies have shown that T-cell responses, particularly against internal viral proteins, are more conserved and correlate with reduced disease severity and cross-variant recognition. Epitope prediction algorithms and immunoinformatics approaches have been widely used to nominate candidate T-cell targets; however, systematic functional validation of conserved non-spike antigens across betacoronaviruses in primary human immune systems, combined with antigen presentation data and *in vivo* vaccine testing, has remained limited. Searches of PubMed and bioRxiv up to December 2025 using terms including “coronavirus T-cell vaccine,” “conserved coronavirus epitopes,” “betacoronavirus cross-reactive T cells,” and “mRNA T-cell vaccine” identified studies demonstrating cross-reactive T-cell immunity and computational epitope selection, but few integrated machine-learning–guided antigen prioritization with *ex vivo* human functional screening, immunopeptidomics, and *in vivo* mRNA immunization in a unified workflow.

*Added value of this study:* This study provides an integrated experimental and computational framework for identifying and validating conserved non-spike T-cell antigens across betacoronaviruses. We functionally screened a panel of candidate peptides derived from multiple viral proteins and demonstrated that evolutionary conservation across species is strongly associated with T-cell immunogenicity. We further demonstrate that multiepitope mRNA constructs encoding these top-ranked conserved regions can be intracellularly expressed, presented on HLA class I molecules to induce polyfunctional T-cell responses in primary human PBMCs. Finally, *in vivo* mRNA immunization in mice induces robust interferon-γ and interleukin-2 T-cell responses exceeding those induced by unconserved control constructs. Together, these findings link evolutionary conservation to functional cellular immunogenicity and extend beyond in silico prediction by demonstrating antigen processing, presentation, and immunogenicity across human and murine systems.

*Implications of all the available evidence:* Collectively, the available evidence indicates that T-cell immunity directed toward conserved internal coronavirus proteins represents a complementary and potentially more variant-tolerant axis of vaccine design than spike-only strategies. Our findings suggest that evolutionary conservation can serve as a practical selection principle for prioritizing T-cell antigens with broad lineage coverage and that multiepitope mRNA delivery is a feasible platform for inducing such responses. While direct protection and heterologous challenge studies will be required to establish clinical efficacy, the integration of computational prioritization with functional validation supports a scalable approach to pandemic preparedness that may be applicable to other rapidly evolving viral families.

## Introduction

Coronaviruses (CoVs) are enveloped, positive-sense RNA viruses classified into four genera (α-, β-, γ-, and δ-CoV). Among alphacoronaviruses (α-CoVs), HCoV-229E and HCoV-NL63 circulate endemically in humans and are typically associated with mild respiratory disease. In contrast, the betacoronavirus (βCoV) genus comprises multiple phylogenetic lineages with distinct ecological niches and zoonotic potential [1, 2], encompassing viruses responsible for disease ranging from endemic respiratory infections to highly pathogenic outbreaks such as SARS-CoV (2003), MERS-CoV (2012), and SARS-CoV-2 (2019) [3]. β-CoVs are further subdivided into five phylogenetic lineages (A–E), corresponding to the subgenera Embecovirus (A), Sarbecovirus (B), Merbecovirus (C), Nobecovirus (D), and Hibecovirus (E) [4], each with characteristic host ranges and pathogenic profiles [5]. Because of this ecological and genetic diversity, β-CoVs have caused past human outbreaks and include credible candidates for future zoonotic spillover, underscoring the need for vaccines capable of preventing severe disease and reducing transmission [6].

The COVID-19 pandemic provided a unique opportunity to study response and adaptation of the population to infection and vaccination. Several studies have investigated the roots of vaccine response heterogeneity, underscoring the importance of conserved epitopes in shaping protection [7]. Endemic CoV infection is associated with less severe COVID-19 [8], and heterotypic immunity from prior SARS-CoV-2 infection, but not vaccination, correlates with reduced endemic CoV incidence, suggesting contributions from non-spike antigens [9]. Individuals with prior SARS-CoV-1 infection from the 2002–2003 epidemic who were later vaccinated with BNT162b2 have been reported to generate broadly neutralizing, pan-sarbecovirus antibodies [10]. SARS-CoV-1 spike vaccination protects animals from SARS-CoV-2 challenge, likely via conserved epitopes, and cellular and humoral immunity to SARS-CoV-2 have been detected in infection-naïve individuals [11–16]. Moreover, prior exposures to endemic β-CoVs have been identified as key factors shaping vaccine outcomes [17–19] and inactivated SARS-CoV-2 vaccines provide partial protection against certain bat sarbecoviruses [20, 21]. Non-structural regions are generally less likely to be exposed and perform essential viral functions, making them less subjected to immune selection pressure. Healthcare workers showed abortive infection with stronger and broader memory T-cell responses than pre-pandemic controls, with preferential recognition of non-structural antigens typical of established infection [22]. These observations have helped delineate strategies for the development of vaccines with broader and more durable protection, making non-structural regions attractive targets for long-term pandemic preparedness [23, 24].

In this study, we combined machine learning–guided antigen prioritization with functional immunological validation to systematically assess conserved CoV T-cell targets across experimental systems. Using a human *ex vivo* vaccination platform, we evaluated the immunogenicity of peptide sequences and mRNA-encoded multiepitope constructs selected for high predicted presentation and evolutionary conservation. To extend these findings beyond in vitro antigen presentation and recall responses, we further assessed the immunogenicity of selected mRNA constructs *in vivo* following vaccination of HLA-A2.1 transgenic mice. Across these complementary models, we examined how sequence conservation relates to antigen processing, TCR activation, and polyfunctionality. Together, this integrated approach enabled direct comparison of peptide immunogenicity and mRNA-based antigen immunization and provided a framework for evaluating conserved epitope-based T-cell vaccine candidates prior to efficacy testing.

## Methods

### Donors

*Ex vivo* validation of T-cell reactivity was performed using PBMCs isolated from healthy donors who had received at least two doses of an mRNA SARS-CoV-2 vaccine (Moderna/mRNA-1273 or Pfizer/BioNTech BNT162b2) and had experienced SARS-CoV-2 infection. PBMCs used in the study were collected between February 11, 2022, and October 11, 2023. Written informed consent was obtained from all participants, and the study was approved by the South-East Norway Regional Ethics Committee.

### Reagents

Commercial spike peptide pools included the PepTivator® SARS-CoV-2 Prot_S Complete (Miltenyi, #130-127-953), consisting of lyophilized 15-mer peptides with 11-amino-acid overlaps covering the full spike glycoprotein (aa 5–1273; Protein QHD43416.1, GenBank MN908947.3). Synthetic peptides were produced by GenScript (Piscataway, NJ, USA) at ≥85% purity, solubilized at 1·5 mg/mL, and stored under standard conditions. All peptides used in the study, including sequence length and protein origin, are listed in **Supplementary Table S1**. PBMC expansion medium consisted of RPMI 1640 with GlutaMAX™ (Thermo Fisher, #61870-010), supplemented with 1 mmol/L sodium pyruvate (Gibco #11360-039), 1 mmol/L MEM NEAA (Gibco #11140-035), 50 nmol/L 1-thioglycerol (Sigma-Aldrich #M1753), 12 µg/mL gentamycin (VWR #E737), and 10% heat-inactivated fetal bovine serum (Gibco #10270-106). IL-2 (R&D #202-IL) and IL-7 (Gibco #PHC0075) were added at 20 U/mL and 1 ng/mL as indicated. Re-challenge assays were performed in TexMACS medium (Miltenyi #130-096-197) supplemented with sodium pyruvate, MEM NEAA, 1-thioglycerol, gentamycin, and 20U/mL IL-2 (R&D #AFL202).

### AI-based Antigen Presentation, Immunogenicity Prediction, and Peptide Selection

All available β-CoV entries were downloaded from BV-BRC, NCBI Virus and UniProt (November 2022). The datasets were then curated to remove sequences with aberrant lengths (i.e., unusually short or long compared to expected protein lengths reported in the literature), as well as incomplete or clearly misannotated entries. Original database annotations were preserved, including accession IDs, source database, organism name, and taxonomic classification, to allow full traceability of each sequence. Next, peptide presentation and immunogenicity of the β-CoV proteome were predicted using the NEC Immune Profiler (NIP) [25, 26], a holistic AI-based platform designed to model the biological processes that lead to antigen presentation on the surface of infected cells and subsequent immune recognition. The platform integrates a series of algorithms that model antigen processing from full-length protein amino acid sequences, including proteolytic processing and intracellular handling, followed by prediction of presentation and peptide-HLA binding affinity, expressed as IC50 values (nM), [25, 26]. The outputs of these AI-modelled biological processes individually reflect the likelihood that a given peptide is processed, presented and bound to a specific HLA molecule on the cell surface. Subsequently, these individual likelihoods are combined into a single metric denoted as antigen presentation (AP) score which ranges from 0 to 1, with 1 being the most likely to be presented on the infected cell surface, [25, 26]. AP score predictions were generated for all possible 9-mer and 10-mer peptides encoded by β-CoV proteome across a panel of 156 most frequent HLA class I alleles in the human population. For each protein and HLA allele considered, a positional AP score was computed for every amino acid, reflecting the likelihood that a minimal epitope encompassing that position would elicit an immune response. These positional scores were derived by assessing immunogenic potential through the AP scores of all overlapping 9-mer and 10-mer peptides containing a given position. Next, immunogenic “hotspots” at protein level were identified by aggregating the AP scores of each amino acid along each protein sequence across all considered HLA class I alleles. A hotspot region was defined as a contiguous protein region with consistently higher AP scores than the rest of the protein sequence. To identify epitopes that are both predicted to be presented and conserved across species, the identified hotspots were clustered based on sequence similarity and amino acid composition. Within each cluster, highly conserved regions were identified as sub-regions of the hotspots that are identical or show very little variation in their amino acid sequence across the entire set of hotspots within that cluster. Each conserved region within a cluster was assigned a conservation score [25, 26], relative to the representative conserved region of that cluster (i.e. the most prevalent conserved region within a cluster). The average AP score of each conserved region across all its constituting amino acids was also calculated. These resulting peptides served as input in the identification of the potential vaccine element candidates. As a result, peptides located within conservation-filtered hotspots and meeting AP > 0·7 and IP > 0·6 for ≥1 Norwegian allele were selected for validation using 15 class I HLA alleles common in the Norwegian population.

To create vaccine constructs, a multi-objective optimization algorithm was used to select the optimal candidate epitope sets from the identified conserved regions, that balance high antigen presentation likelihood, conservation across species and broad HLA coverage, thereby enabling the design of vaccine candidates with potential cross-taxon protective capacity. This approach guided the selection toward vaccine designs that maximize the strain and species coverage while maintaining high AP scores and broad HLA reactivity. As a result, two vaccine constructs, namely NEC-T4, NEC-T5, and NEC-T6 were made, including 11,10, and 20 hotspots, respectively (**Supplementary Table S3**).

### mRNA Construct Design, Synthesis and formulation

Three mRNA vaccine constructs, NEC-T4, NEC-T5, and NEC-T6, were designed to encode concatenated T-cell antigenic regions predominantly derived from non-structural CoV proteins (**Supplementary Table S1**). Antigenic regions were selected based on predicted immunogenicity, conservation, and HLA coverage, and were arranged as poly-epitope sequences within a single open reading frame. The constructs incorporated an N-terminal signal sequence followed by a dendritic cell lysosomal-associated membrane protein (DC-LAMP) targeting domain to promote intracellular trafficking of the translated poly-epitope protein to the endosomal/lysosomal compartment [27]. This design facilitates antigen processing and presentation through the major histocompatibility complex (MHC) pathways, thereby enhancing T-cell priming. Plasmid production, mRNA synthesis, and formulation were performed by eTheRNA immunotherapies NV (Niel, Belgium) using their proprietary mRNA vector platform. DNA templates for *in vitro* transcription were generated from plasmids encoding the full vaccine constructs, including the signal sequence and DC-LAMP domain. mRNA was synthesized by *in vitro* transcription using wild-type nucleoside triphosphates and incorporated a co-transcriptional 5′ cap structure using the CleanCap technology to ensure efficient translation. A co-transcriptional poly(A) tail was included to enhance mRNA stability and translational efficiency. Following transcription, mRNA was purified according to eTheRNA’s established protocols and formulated for downstream experimental use. All steps of mRNA production and formulation were conducted under conditions optimized to preserve RNA integrity, translational competence, and immunogenic performance.

### PBMC Isolation and Biobanking

PBMCs were isolated using CPT tubes (BD Vacutainer #362782) according to the manufacturer’s protocol. After centrifugation (1,600 g, 25 min, RT), PBMCs were washed in cold PBS (Gibco #10010-015), counted, and resuspended in FBS containing 10% DMSO. Aliquots of ≥5 × 10L cells per vial were frozen overnight at −80 °C in Mr. Frosty containers (Nalgene #5100-0001) and transferred to liquid nitrogen for long-term storage.

### PBMC *Ex Vivo* Vaccination

PBMCs were thawed, and dead cells were removed (Dead Cell Removal Kit, Miltenyi Biotec). Viable cells were counted and split into two equal fractions (1 × 10L cells each), plated in 48-well plates (0·5 mL/well), and rested for ≥4 h. For peptide-based stimulation, one fraction was incubated for 1 h with peptide pools B1, B2, or B3 (2 µg/mL per peptide) or Peptivator (0.6 nmol) and then combined with the untreated fraction. For mRNA construct-based stimulation, one fraction was irradiated, washed, and incubated for 24 h with LNP-formulated NEC-T4 or NEC-T5 (0·1 to 10 µg/mL). Cells were washed and mixed with untreated PBMCs.

Cells were expanded with half-media changes every 3–4 days using medium supplemented with IL-2 (20 U/mL) and IL-7 (1 ng/mL). Restimulation occurred on days 15 and 25 using fresh irradiated peptide- or construct-loaded autologous PBMCs (3 × 10L cells in 300 µL). On day 35, cells were harvested for rechallenge-induced T-cell activation.

### T-cell Activation-Induced Marker (AIM) Assay and Flow Cytometry

PBMCs (1 × 10L cells/mL) were resuspended in TexMACS medium (200 µL/well) and incubated for 3 h with Peptivator (0·6 nmol) or peptide pools (B1, B2, or B3) at a final concentration of 1·5 µg/mL. per peptide for construct testing. PBMCs were stimulated with autologous PBMCs incubated overnight with either NEC-T4 or NEC-T5. After overnight incubation in presence of Brefeldin A/Monensin (GolgiStop, Invitrogen #00-4980-93) cells were washed in cold PBS and processed for flow cytometry. Cells were washed in FACS buffer (PBS + 1% BSA) and stained with Fixable Near-IR Live/Dead dye (Molecular Probes #L34976) for 10 min. After surface staining with BV605 anti-CD3 (BD #563219), PerCP-Cy5.5 anti-CD4 (BioLegend #317428), and Alexa Fluor 488 anti-CD8 (Invitrogen #53-0086-42), cells were permeabilized using BD Perm/Wash, stained with APC anti-CD137 (BD #550890), BV711 anti-CD40L (BioLegend #310837), PE anti-IFN-γ (BioLegend #502509), and BV421 anti-TNF (BD #562783), washed twice in FACS buffer, and acquired on an Attune NxT Flow Cytometer (Thermo Fisher).

### Reactivity Score and Polyfunctionality

T-cell reactivity scores (RS) for CTL and Th subsets were calculated as previously described [25, 28]. Briefly, RS values were derived from activation-induced marker expression across predefined T cell populations and normalized to background-subtracted responses. For each T-cell subset, RS was computed by averaging normalized responses from selected informative populations, with negative values set to 0·0001. Polyfunctional profiling was performed using FlowJo 10.10 Boolean analysis and visualized using Simplified Presentation of Incredible Complex Evaluations (SPICE) 6.1 [29–31].

### Global Proteomics and LC–MS Analysis

#### Cell culture and transfection

Cell culture and transfection were carried out as follows. Expi293F cells (Thermo Fisher Scientific), a HEK293-derivative that expresses HLA A*02:01, A*03:01 and HLA-B*07:02 (Cellosaurus), were maintained in suspension in serum-free medium according to the manufacturer’s instructions. For initial optimization, cells were transfected with GFP lipid nanoparticles (LNPs) at 500 ng/mL. For assay setup, 300,000 cells per well were seeded in complete DMEM supplemented with 10% fetal bovine serum (FBS). Transfection efficiency was assessed 20–24 h later by fluorescence microscopy and flow cytometry. For experimental transfections, 75 µg of each LNP formulation was diluted in 75 mL of complete medium and combined with 75 mL of Expi293F cells at a density of 600,000 cells/mL. The resulting suspension was divided equally into three T175 flasks corresponding to sampling time points of 6, 24, and 48 h. Cultures were incubated at 37 °C in a humidified atmosphere containing 5% CO₂. At each time point, cells were harvested using Accutase (Innovative Cell Technologies), washed with phosphate-buffered saline (PBS), snap-frozen on dry ice, and stored at –80 °C until proteomic analysis.

#### Protein extraction and digestion

Frozen cell pellets were washed three times with PBS and lysed in 8 M urea, 50 mM Tris-HCl (pH 8·0), supplemented with Complete Protease Inhibitor (MilliporeSigma). Lysates were sonicated at 35–40% amplitude (1s on/1s off pulses, 20 s total) and clarified by centrifugation at 14,000 × g for 10 min at 4 °C. Protein concentrations were determined using a Qubit assay (Thermo Fisher Scientific). For digestion, 50 µg of protein from each sample was reduced with 12 mM dithiothreitol (DTT) for 1 h at room temperature and alkylated with 15 mM iodoacetamide for 1 h in the dark. Proteins were digested overnight (18 h) at 37 °C with sequencing-grade trypsin (Promega) at a 1:20 (w/w) enzyme-to-substrate ratio. Digests were acidified to a final concentration of 0·3% trifluoroacetic acid (TFA) prior to solid-phase extraction (SPE). Peptides were purified using Oasis µHLB SPE plates (Waters). The matrix was activated with four washes of 70% acetonitrile (500 µL each) and equilibrated with four washes of 0·3% TFA. Samples were loaded, washed three times with 0·3% TFA, and eluted with 200 µL followed by 400 µL of 60% acetonitrile/0·3% TFA. Combined eluates were frozen at –80 °C and lyophilized overnight.

#### LC–MS/MS and data analysis

Dried peptides (500 ng per injection) were analyzed on a Vanquish Neo UPLC system (Thermo Fisher Scientific) coupled to an Orbitrap Astral mass spectrometer (Thermo Fisher Scientific), operated in DIA mode with MS1 spectra acquired in the Orbitrap analyzer and MS2 spectra acquired in the Astral analyzer. Peptides were loaded onto a trapping column and separated on a 75 µm analytical column maintained at 55 °C using a 30-min gradient at a flow rate of 350 nL/min. The mass spectrometer was operated in data-independent acquisition (DIA) mode. Full MS scans were acquired at a resolution of 240,000 (FWHM) over an m/z range of 380–980. DIA scans employed 300 × 2 m/z isolation windows, with fragment ions detected at 40,000 FWHM in the Astral analyzer. The normalized collision energy (NCE) was set to 25, and the maximum ion injection time was 3·5 ms. A pooled quality control (QC) sample was injected at the beginning and end of each batch. DIA data were processed using DIA-NN (v1.9.1). Raw files were converted to QUANT outputs, retention times were aligned, and spectra were searched against an in silico spectral library generated from a FASTA database. An iterative search was subsequently performed using an empirical spectral library generated by DIA-NN. Data were filtered at a 1% peptide- and protein-level false discovery rate (FDR). Peptide intensities were quantified as area-under-the-curve values and normalized across samples.

### Immunopeptidomics

Expi293F suspension cells were expanded in Expi293 medium to a total of over 5×10^8^ cells. Cells were then resuspended in complete medium consisting of DMEM supplemented with 10% FBS and penicillin/streptomycin at a density of 6×10^5^ cells/mL. For each of the six LNP formulations, 100 µg of LNP-encapsulated mRNA was diluted in 100 mL of complete medium and mixed with 100 mL of Expi293F cell suspension in sterile bottles. Each transfection mixture was transferred to Falcon 5-layer tissue culture treated flasks (Corning). As a negative control, 200 mL of cells at 3×10^5^ cells/mL were seeded into a 5-layer flask without LNP treatment. In parallel, GFP-encoding LNPs were transfected into Expi293F cells using the same procedure and seeded into T25 flasks to assess transfection efficiency by flow cytometry prior to harvesting. Cells were incubated at 37°C with 5% CO₂ for 48 hours. Following incubation, cells grown in 5-layer flasks were gently rinsed with PBS and detached using Accutase (Gibco). Cells were pelleted by centrifugation at 300×g for 15 minutes with slow deceleration. Pellets were washed once with 25 mL PBS, counted, and centrifuged again at 300×g. Cell pellets were snap-frozen on dry ice and stored at −80°C until further processing. Frozen pellets were thawed on ice and lysed at a concentration of 5×10^7^ cells/mL in lysis buffer, followed by incubation on ice for 30 minutes. Insoluble material was removed by centrifugation at 800×g for 5 minutes at 4°C. The resulting supernatant was further clarified by centrifugation at 20,000×g for 30 minutes at 4°C. Clarified cell lysates were incubated with pre-equilibrated anti-HLA class I (W6/32) immunoaffinity resin under gentle rotation at 4°C overnight to capture HLA–peptide complexes. The following day, resin was pelleted by centrifugation at 800×g for 5 minutes at 4°C. Three sequential washes were performed using wash buffers 1–3. For each wash, 2·5 mL of buffer was added, samples were vortexed, centrifuged at 800×g for 5 minutes at 4°C, and the supernatant discarded. A fourth wash was performed by adding 0·75 mL of wash buffer 4, after which the resin was transferred to LoBind tubes (Eppendorf) and centrifuged again at 800×g for 5 minutes at 4°C. Supernatants were discarded. Bound peptides were eluted by incubating the resin with 1 mL of elution buffer at 37°C for 5 minutes, followed by centrifugation at 800×g for 5 minutes at 4°C. Eluates were collected into fresh LoBind tubes and stored at −80°C until mass spectrometry analysis. Half of each eluate was desalted using solid-phase extraction (SPE) with a µHLB C18 plate (Waters). Peptides were loaded directly onto the SPE material and eluted using 30/70 (v/v) acetonitrile/water containing 0·1% trifluoroacetic acid (TFA). Eluted peptides were lyophilized and reconstituted in 0·1% acid prior to analysis. For comprehensive analysis, 100% of the peptide material was analyzed by nano–liquid chromatography tandem mass spectrometry (nanoLC–MS/MS) using a Waters NanoAcquity system interfaced to a Thermo Fisher Scientific Fusion Lumos mass spectrometer. Peptides were first loaded onto a trapping column and subsequently separated on a 75 µm analytical column at a flow rate of 350 nL/min. Both columns were packed with Luna C18 resin (Phenomenex). A 2-hour linear gradient was employed for peptide separation. The mass spectrometer was operated in data-dependent acquisition mode. Full MS scans were acquired in the Orbitrap at a resolution of 60,000 (FWHM), followed by sequential MS/MS scans using high-resolution CID and EThcD in the Orbitrap at 15,000 (FWHM). Spectra were acquired over an m/z range of 300–1600, with a cycle time of 3 seconds. Raw mass spectrometry files were searched using a local installation of PEAKS software (Bioinformatics Solutions Inc) against the SwissProt human protein database supplemented with custom sequences. Searches were performed with no enzyme specificity. Variable modifications included methionine oxidation, protein N-terminal acetylation, and carbamidomethylation of cysteine. Monoisotopic mass values were used with a precursor mass tolerance of 10 ppm and a fragment mass tolerance of 0·02 Da. Peptide-spectrum matches were filtered to a false discovery rate (FDR) of 1%. Chimeric peptide identification was enabled.

### Discriminatory performance of the immunogenicity score

The discriminatory performance of the immunogenicity score was evaluated using receiver operating characteristic (ROC) curve analysis and area under the curve (AUC) metrics. Percentile rank scores were calculated for each peptide–HLA pair (lower values indicate stronger predicted immunogenicity) and used solely for visualization purposes. Analyses were performed for vaccine-aligned peptides predicted to be presented by the class I HLA alleles HLA-A02:01, HLA-A03:01, and HLA-B*07:02 of Expi293F cells. Predicted epitopes were first restricted to peptides aligned to the vaccine constructs and to the selected HLA alleles. For each unique peptide–HLA–species combination, only the maximum predicted immunogenicity score was retained, ensuring that each peptide–allele pair contributed a single representative score to the analysis. Peptides identified by immunopeptidomics were designated as positives, while peptides not detected experimentally were designated as negatives. ROC curves were generated by thresholding the continuous immunogenicity score and calculating true positive and false positive rates across all thresholds. AUC values were computed to quantify the ability of the immunogenicity score to discriminate between experimentally detected and non-detected peptides, independent of score threshold. ROC curves were visualized with the false positive rate on the x-axis and the true positive rate on the y-axis, with a diagonal reference indicating random classification performance. All statistical analyses were conducted in Python using standard scientific computing libraries.

### Mice

Thirty-six female HLA-A2.1 transgenic mice (Taconic model 9659; CB6F1-Map4k3^Tg(HLA-A0201/H2-Kb)A0201^/Tac), aged ∼8 weeks were randomized into six groups (n = 6 each): 1) Comirnarty reference vaccine, 2) GFP-LNP control, 3) full length spike-LNP (Wuhan), 4) Construct-NEC-T4, 5) Construct NEC-T5, or 6) unvaccinated controls. Mice received intramuscular injections (50 µL into the hind limb) on days 0, 21, and 42 with 10 µg of mRNA–LNP or 1 µg of Comirnarty. All procedures complied with institutional and national ethical regulations. Spleens were collected aseptically on days 54 and 56, mechanically dissociated through 100 µm strainers into ice-cold HBSS with 1.5% FBS and 1% penicillin–streptomycin and subjected to erythrocyte lysis (1× eBioscience RBC lysis buffer). Cells were washed, passed through 70 µm strainers, and resuspended in complete RPMI-1640 (10% FBS, 2 mM L-glutamine, 1% penicillin–streptomycin, 0·05 mM 2-mercaptoethanol). Viable cell numbers were determined by trypan blue exclusion (Bio-Rad TC20). Splenocytes were rested for ∼1 h at 37 °C, 5% CO₂ prior to stimulation. For antigen re-challenge, cells were stimulated with overlapping 15-mer peptide pools and analyzed using triple-color FluoroSpot (IFN-γ, IL-2, TNF-α; Mabtech #FSP-414245). Custom peptides (≥75% purity; GenScript PepHTS™) consisted of 298 overlapping 15-mers (12 aa overlap) spanning the mRNA constructs and organized into subpools OVL-4A/B (NEC-T4) and OVL-5A/B (NEC-T5). Lyophilized peptides were stored at –20 °C, reconstituted in DMSO, and used at 1 µg/mL per peptide (≤0·25% final DMSO). Additional stimulations included spike’s 15-mer pools (Pepscan) and DMSO controls; Concanavalin A served as a positive control. Anti-CD28 (0·2 µg/mL) was added to all wells. Splenocytes (2·5 × 10L per well) were plated in duplicate on pre-coated FluoroSpot plates, blocked with complete medium, and incubated for 18–19 h at 37 °C, 5% CO₂. Reference splenocytes from Comirnaty-vaccinated BALB/c mice were included for inter-assay calibration. Following incubation, plates were washed and sequentially developed with biotinylated detection antibodies and fluorophore-conjugated secondary reagents (FITC for IFN-γ, Cy3 for IL-2, Cy5 for TNF), enhanced, air-dried, and imaged within 72 h using an AID vSpot Spectrum reader. Spot quantification was performed using AID EliSpot v7.0 with standardized exposure settings and automated gating without manual adjustments. Single-, double-, and triple-cytokine spot forming units (SFUs) were calculated according to the manufacturer’s guidelines and normalized to 100%, with no background subtraction. Duplicate wells were averaged for each animal and condition; data from Days 56 and 57 were combined. Plate-level QC confirmed consistent calibration across assays.

### Statistics

Statistical significance was assessed using Mann–Whitney U tests, Fisher’s exact tests, one-way ANOVA, or Friedman’s test, with Dunnett’s multiple comparisons test applied as appropriate using GraphPad Prism v10.

## Results

### Identification of conserved epitopes for T-cell response analyses

All available β-CoV protein sequences (*n* = 8942750) were compiled and curated to generate a non-redundant reference proteome (*n* = 4000; see Methods). Antigen presentation and T-cell response likelihood was then predicted across the β-CoV proteome using the NEC Immune Profiler, yielding antigen presentation (AP) scores for all overlapping 9- and 10-mer peptides across a broad panel of human HLA class I alleles (**Supplementary Table S1**, columns AF-ME). Aggregation of positional AP scores identified discrete immunogenic hotspots within individual viral proteins. These hotspots were subsequently clustered by sequence similarity to define conserved regions shared across multiple CoV taxa (see Methods). Conserved hotspot regions with high predicted likelihood of T-cell response were prioritized and used as input for vaccine design, resulting in the selection of multiepitope mRNA constructs optimized for both high AP scores and broad taxonomic coverage. The β-CoV genera includes the subgenera sarbecovirus, embecovirus, merbecovirus, nobecovirus, and hibecovirus, as well as unclassified βCoV. In **Supplementary Figure S1A** we show subgenera vs distinct CoV taxa (defined as individual viral isolates or reference sequences) for the peptides included in the analysis and listed in **Supplementary Table S1**. Most peptides were present in 2–6 subgenera, with the majority also conserved across 129–289 distinct taxa, defined as individual viral isolates or reference sequences. Of the 205 peptides analyzed, 89 were conserved exclusively within Sarbecovirus, while 86 extended to closely related unclassified β-CoVs (**Supplementary Figure S1B**). A smaller subset of 30 peptides showed broader conservation across multiple β-CoV subgenera, including 3 peptides conserved across all six recognized β-CoV subgenera (**Supplementary Figure S1B**). Within the sarbecovirus subgenera, most peptides (*n* = 142) were shared across SARS-CoV, bat COV and pangolin COV taxa (**Supplementary Figure S1C)**.

### Degree of epitope sharing is associated with higher reactivity and response frequency

To assess whether broad sequence conservation is associated with T-cell immunogenicity, we first analyzed previous response data from 97 SARS-CoV-2 peptides for which reactivity data were available [25]. These SARS-CoV-2 peptides (**Supplementary Table S1,** Federico et al., column G) were either private for SARS-CoV-2 or conserved across distinct CoV taxa spanning multiple host species (**Supplementary Table S1,** columns S and U). We applied an arbitrary threshold based on the number of distinct CoV taxa across which each peptide was conserved (**Supplementary Table S1**, column U), classifying peptides as conserved when found in more than five taxa. Using this stratification, we observed that low stimulatory peptides defined by a Reactivity Score [RS] ≤ 0·0001 (see **Methods**) were predominantly enriched among peptides conserved across five or fewer taxa (**Supplementary Figure S2**,). This pattern was evident in both CTL and Th compartments. In contrast, peptides conserved across more than five taxa were more likely to elicit measurable T-cell responses, To quantify these differences, we compared mean reactivity scores across donors. Peptides with broader sequence conservation across CoV taxa (>5) exhibited significantly higher reactivity than less conserved peptides (≤ 5) for both CTLs and Th cells (median relative reactivity: 2·6 vs. 0·0001 for CTL; 1·1 vs. 0·0001 for Th; p = 0·0071, Mann–Whitney test; **Figure 1A**). This distinction was also evident at the individual donor level: with the exception of for donors 114 and 117 (both CTL and Th responses) and donor 112 (only CTL), all donors showed greater reactivity to peptides with broader taxonomic conservation (**Figure 1B**). Consistent with these findings, the proportion of reactive peptides was higher among broadly conserved epitopes (72% vs. 30% for CTL; 46% vs. 22% for Th; p < 0·001, Fisher’s exact test; **Figure 1C**). This pattern was largely maintained in per-donor analyses, with the exception of donors 110, 112, and 131 within the CTL compartment (**Figure 1D**). Notably, 81% of antigenic sequences with broad conservation mapped to non-structural regions (Orf proteins), while only 19% mapped to structural regions (spike and E proteins), respectively. Collectively, these results indicated that sequence conservation across diverse CoV taxa, spanning multiple viral isolates and host origins, was strongly associated with increased likelihood and magnitude of T-cell recognition across individuals

**Figure 1.**
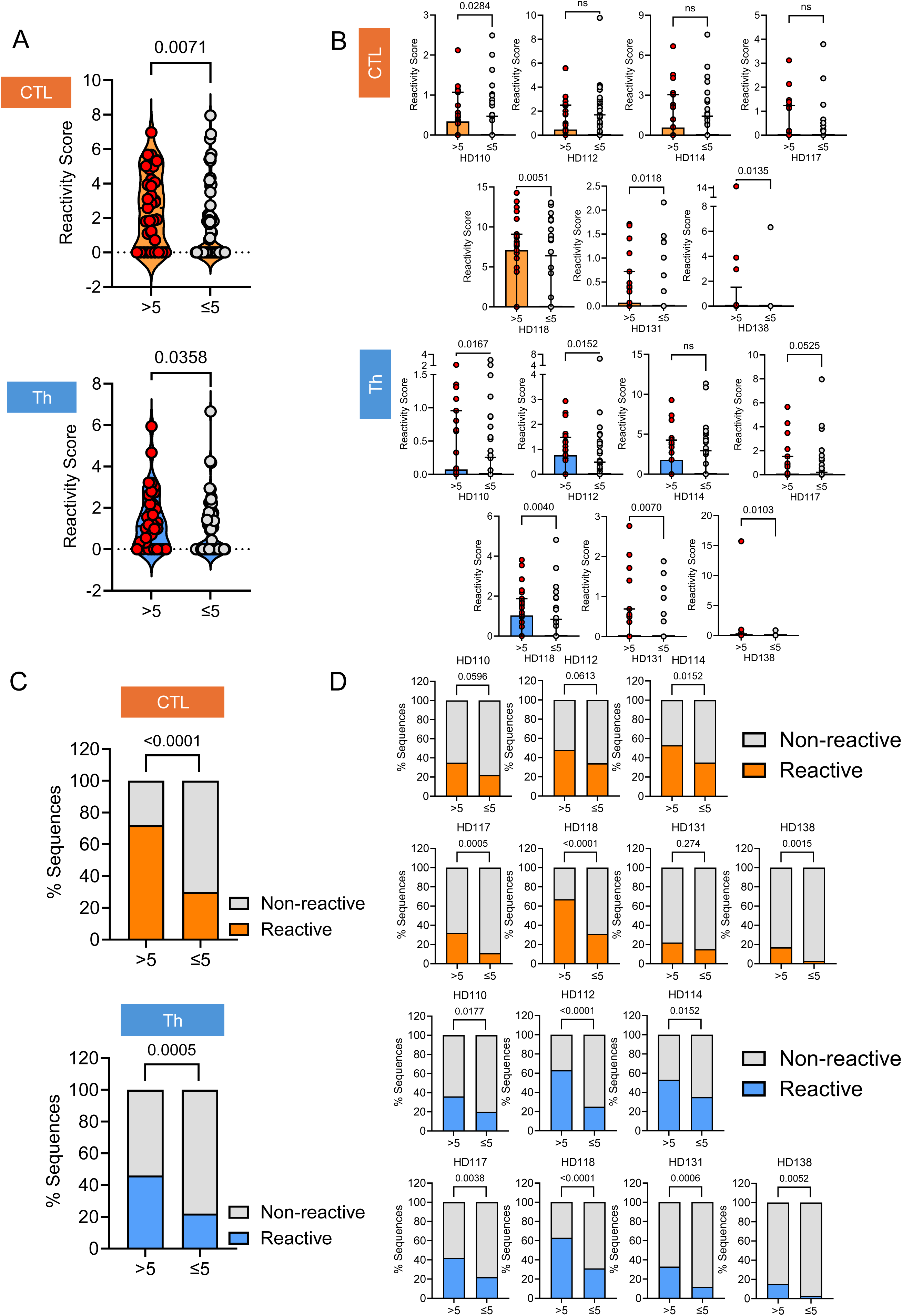
Degree of epitope sharing is associated with higher reactivity and response frequency. (**A**) Average reactivity scores across all donors (*n* = 7) for individual peptides, stratified by degree of sharing (cut-off: ≤5 species). (**B**) Distribution of reactivity scores per donor. (**C–D**) Percentage of reactive peptides in CTL and Th compartments, stratified by degree of sharing (C) and per donor (D). Statistical comparisons were performed using the Mann–Whitney U test (A, B) and Fisher’s exact test (C, D). Significance: *p < 0·05; **p < 0·01.

### Response to *ex vivo* vaccination with SARS-CoV-2 peptides is detectable and specific

To rapidly assess candidate conserved β-CoV antigens we implemented and validated a T-cell stimulation workflow—hereafter referred to as the *ex vivo* vaccination procedure (see methods)—to determine whether stimulus-specific T cells could be selectively expanded from PBMCs of healthy vaccinated donors. PBMCs were stimulated with either (i) 9–10-mer peptides predicted by the NEC Immune Profiler (NIP) AI platform [25, 26] spanning the spike protein or non-structural proteins, grouped into (i) two heterogeneous pools (B1 and B2), or (ii) Peptivator, a commercially available pool of overlapping 15-mer peptides covering the full spike protein. The genera and subspecies expression, HLA binding and characteristics of the pools are provided in **Supplementary Table S1**. Following the initial challenge, cells underwent two additional stimulations on day 15 (D15) and day 25 (D25) to promote expansion of reactive T-cell clones. Antigen specificity of the resulting CTL and Th lines was assessed after re-challenge at day 35, using flow cytometric analysis of activation-induced markers (AIMs), including CD137, CD40L, and IFN-γ (**Supplementary Figure 3A** and **Methods**). Each stimulus elicited robust and selective expansion of antigen-specific T cells: As shown in **Supplementary Figure 3B**, stimulation with B1, B2 and Peptivator pools produced significant enrichment in reactive cells in the CTL compartment (64% and 66% of B1- and B2-reactive CD137⁺IFN-γ⁺ cells, respectively) and in the Th cell compartment (18% of B1-reactive CD40L⁺CD137⁺ cells). As shown in **Supplementary Figure 3C–D**, enrichment of stimulus-specific T cells occurred to varying degrees in all tested donors, with up to ∼65% of both CTL and Th populations becoming activated upon restimulation: Donor-specific variation was evident. For example, CTLs from donor 124 showed poor expansion following B2 stimulation (**Supplementary Figure 3C**, middle panel), whereas Th cells from the same donor responded strongly to the Peptivator (**Supplementary Figure 3D**, middle panel), likely reflecting sequence overlap among peptide pools. Consistently, the Peptivator pool preferentially expanded Th cells (**Supplementary Figure 3D**, lower panels), in line with the known preference of 15-mer peptides for MHC class II–restricted presentation. In a cohort of 17 healthy donors, the *ex vivo* vaccination procedure consistently supported variable but robust expansion of antigen-specific CTL and Th populations, quantified as the fraction of cells double-positive for CD137, CD40L, IFN-γ, or TNF (**Supplementary Figures 3E-F**). Overall, these data demonstrate that, despite the observed inter-donor heterogeneity, this stimulation protocol generates highly specific antigen-specific T-cell expansion and can be used for *ex vivo* testing of T-cell targets.

### Polyfunctional response to *ex vivo* vaccination with conserved **β**-CoV peptides is detectable in all donors

To assess the frequency of responses to conserved β-CoV sequences, we focused on peptide pools B2 and B3, which were enriched for sequences shared across multiple subgenera and taxa (**Supplementary Table S1**). To evaluate the functional quality of *ex vivo*–expanded T cells, we quantified the frequency of polyfunctional antigen-specific CTL and Th cells. Polyfunctionality was defined as the proportion of cells co-expressing multiple activation-induced markers (AIMs) and/or cytokines (**Supplementary Table S2**), and combinatorial marker expression was visualized using the SPICE analysis tool (**see Methods**). At day 35 post-*ex vivo* expansion, antigen-specific T cells capable of simultaneously expressing multiple activation markers were detected in all donors. Following stimulation with the B2 pool, polyfunctional CTLs and Th cells were detected in all tested donors (*n* = 7), ranging from 2·1% to 84% (CTL; **Figure 2A**) and 3% to 38% (Th; **Figure 2B**) of the activated population, respectively. Only one donor (203) displayed a minimal Th response (0·3%; **Figure 2B**). Further analysis of polyfunctional subsets revealed that four of seven donors exhibited large fractions (>30%) of IFN-γ⁺CD137⁺TNF⁺ triple-positive CTLs (**Figure 2C**; **Supplementary Table S2**). Th polyfunctionality varied more widely, with reduced responses in donors 200 and 204 (**Figure 2D**). Notably, donor 203—despite low overall Th activation—showed a disproportionately high fraction of triple- and quadruple-positive Th cells (∼25% quadruple-positive; **Figure 2D**).

**Figure 2.**
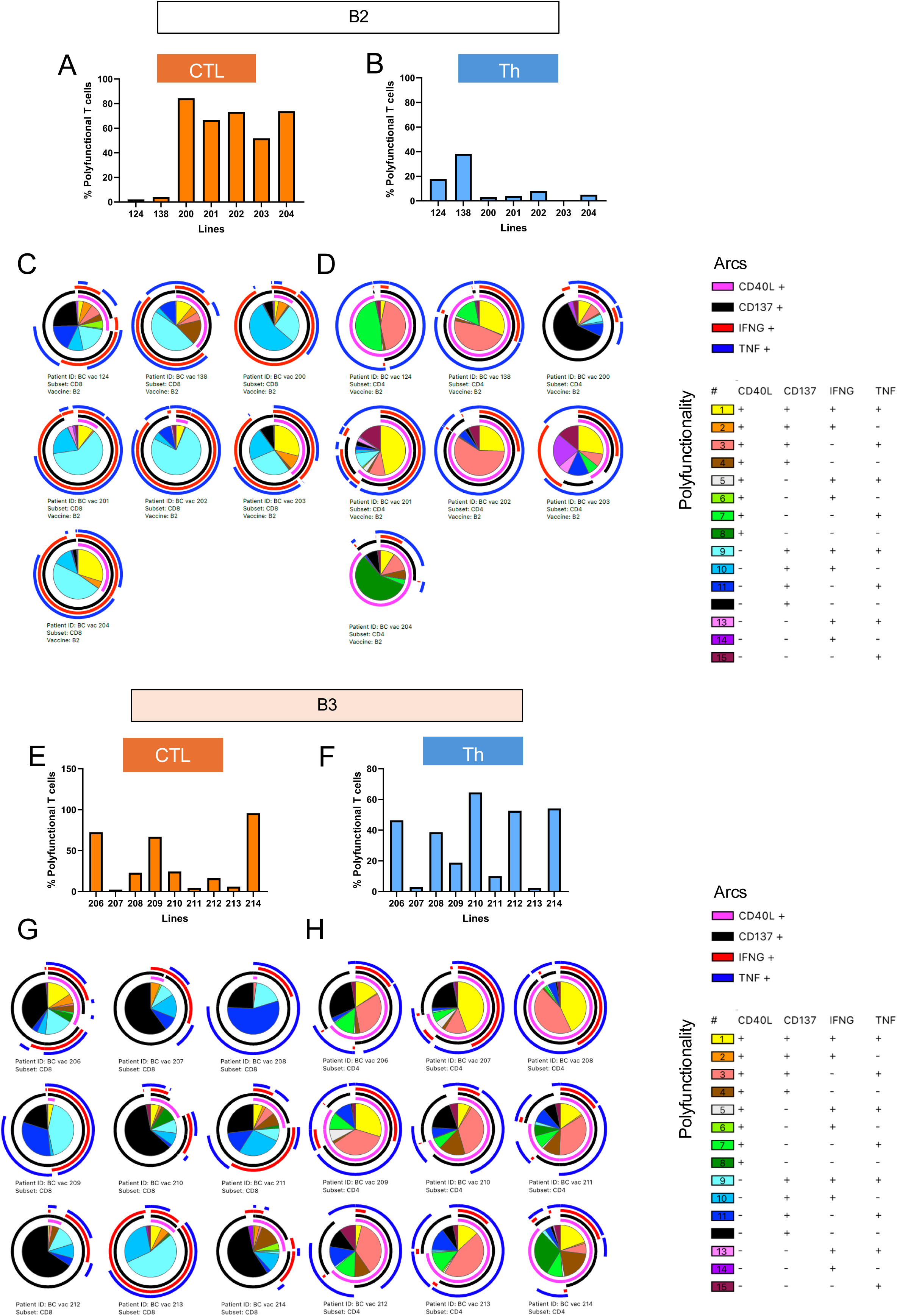
Polyfunctional response to *ex vivo* vaccination with conserved β-CoV peptides is detectable in all donors. (**A-B**) Percentage of polyfunctional CTL (A) and Th (B) T cells after stimulation with the B2 peptide pool at day 35. (**C–D**) Polyfunctional fractions defined by marker co-expression. Overlapping arcs indicate co-expression, and pie charts show the relative abundance of each polyfunctional subset for CTL (C) and Th (D). Pie chart colors indicate the relative abundance of polyfunctional subsets, with color coding defined in the legend. (**E–F**), and (**G-H**) Corresponding analyses for CTL and Th cells after stimulation with the B3 peptide pool.

Stimulation with the B3 pool produced patterns similar to those observed with B2. Multiple donors exhibited detectable polyfunctional responses in both T-cell compartments, and responses were observed in all donors (*n* = 9; **Figure 2E–F**). In the CTL compartment, higher-order polyfunctional subsets were detected (**Figure 2G**), and this trend was even more pronounced in Th cells, which contained larger fractions of triple- and quadruple-positive populations in several donors (**Figure 2H**; **Supplementary Table S2**). Overall, polyfunctional T-cell responses were detected in all donors and across all peptide pools tested. These findings demonstrate that *ex vivo* stimulation with conserved β-CoV peptides consistently induces multifunctional T-cell immunity.

### Design and expression of mRNA-based CoV constructs

To build on our findings regarding the relationship between sequence conservation and antigenicity, we generated three mRNA constructs (NEC-T4, NEC-T5, and NEC-T6) encoding concatenated T-cell antigenic sequences predominantly derived from non-structural CoV proteins (**Supplementary Table S1**). These sequences were selected for broad conservation across β-CoV subgenera—including sarbecoviruses, and other CoVs with zoonotic spillover potential (**Figure 3A**) [32] and included some epitopes present in the B1, B2, and B3 pools, along with additional peptides conserved across CoV species (**Supplementary Table S1**; **Methods**). We first evaluated construct expression in Expi293F cells and monitored protein abundance over time using global proteomics. Vaccine-derived proteins were detectable as early as 6 hours post-transfection and increased steadily, peaking at 24–48 hours (**Figure 3B**). Ranking proteins by mean intensity across the time course showed that for 2 of the 3 constructs (NEC-T4 and NEC-T5) construct-encoded proteins reached expression levels comparable to mid- to high-abundance endogenous proteins (**Figure 3C**), confirming robust expression. To assess antigen processing and presentation through the MHC class I pathway, we performed MHC-I immunoprecipitation on cells harvested 48 hours post-transfection. Peptides derived from these constructs were detected (**Supplementary Figure S4A**), demonstrating active intracellular processing and presentation. We next evaluated the distribution of MHC-presented peptides across CoVs subgenera. Construct-derived peptides exhibited broad coverage across major lineages: identified sequences mapped to five of six β-CoV subgenera, with species coverage ranging from 88% (230/258) to 89% (36/41) per subgenus. Only unclassified β-CoVs showed limited representation (2 of 146 species; **Supplementary Figure S4B**). To assess predictive performance of the immunogenicity score, receiver operating characteristic (ROC) analysis was performed using immunogenicity scores filtered for the HLA alleles in Expi293F cells (HLA-A*02:01, HLA-A*03:01, HLA-B*07:02). High predictive accuracy was observed as measured by an AUC of 0·85 for all HLA combined (**Supplementary Figure S4C**) and 0·88, 0·82, and 0·92 for each of the three HLA examined (**Supplementary Figure S4D**). Together, these data demonstrate that predicted immunoreactive conserved CoV sequences encoded by mRNA constructs are expressed, processed, and presented *in vitro*. Because of the limited expression of the NEC-T6 construct, we focused our analyses on NEC-T4 and NEC-T5. These constructs were designed to include sequences predominantly derived from non-structural CoV proteins (**Supplementary Table S1**) and exhibited broad conservation across β-CoV subgenera (**Figure 3B–D**). The peptide sequences incorporated into the NEC-T4 and NEC-T5 constructs were shared across 2 to 6 multiple subgenera (*n* = 2 - 6) and taxa (**Figure 3D–E**), and were predominantly conserved among SARS-CoV, bat CoV, and pangolin CoV (**Figure 3F–G**). Together, these results provide a proof-of-concept framework for the design and expression of mRNA constructs incorporating AI-predicted conserved regions for use in evaluating T-cell immunogenicity.

**Figure 3.**
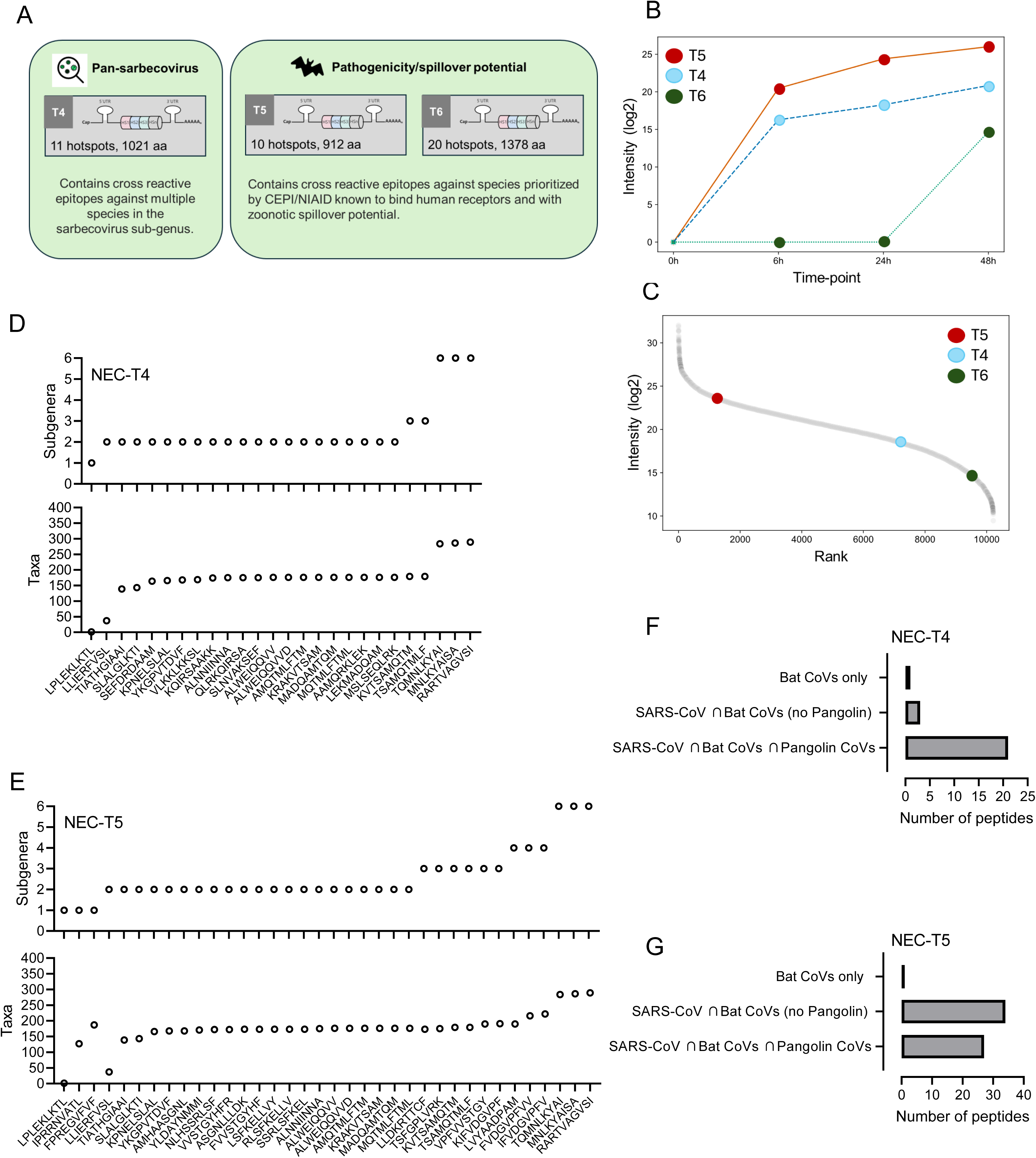
Design and expression of mRNA-based CoV constructs. (**A**) Design of RNA constructs NEC-T4–T6. (**B**) Protein expression kinetics following transfection of Expi293F cells with mRNA constructs at 6, 24, and 48 h. Curves display construct-specific expression trends as log₂ intensity measured by DIA-NN. (**C**) Relative abundance of vaccine-derived proteins compared with the global proteome. Proteins were ranked by mean log₂ intensity across time points; vaccine-derived proteins are highlighted relative to endogenous proteins. (**D-E**). Immunogenic peptides included in the NEC-T4 and NEC-T5 constructs. β-CoV subgenera (top panel) and taxa (bottom panel) counts are shown. (**F-G**) Sarbecovirus-specific conservation. Bars show the number of peptides conserved in each combination of SARS-CoV (SARS-CoV-1 ∪ SARS-CoV-2), bat, and pangolin CoVs.

### *Ex vivo* testing of mRNA-based CoV constructs

We next used our *ex vivo* vaccination platform to assess whether stimulation of PBMCs with mRNA-encoded NEC-T4 and NEC-T5 constructs could expand antigen-specific T-cell. NEC-T6 was not further evaluated because of its suboptimal expression kinetics. Although the magnitude of responses at day 35 following antigenic re-challenge was generally lower than that induced by peptide stimulation (**Figure 2**), we observed a dose-dependent increase in AIM⁺ cells in both the Th and CTL compartments when expanded PBMCs were re-challenged with increasing amounts of construct (1–10 μg/mL), with consistently stronger responses in Th cells. For example, donor 210 (LN210) exhibited up to 16% CD137⁺CD40L⁺ Th cells following re-challenge (**Figure 4A**). In contrast, CTL responses were of lower magnitude, with a peak frequency of 0·6% IFN-γ⁺CD137⁺ cells detected in donor 215 (LN215; **Figure 4B**).

**Figure 4.**
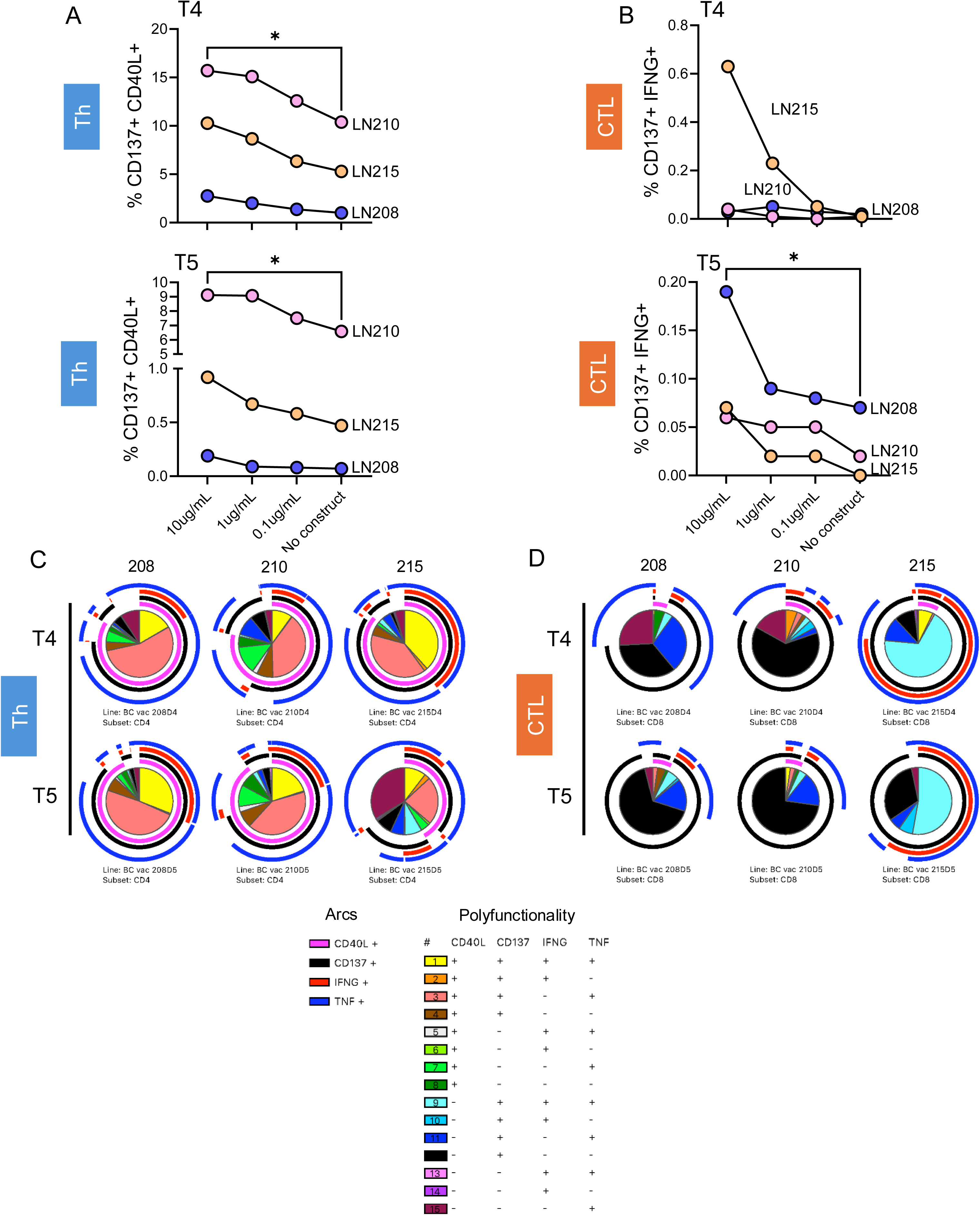
*Ex vivo* testing of mRNA-based CoV constructs. (**A–B**) T-cell activation after stimulation with NEC-T4 or NEC-T5 in the (A) Th and (B) CTL compartments measured on day 35 by co-expression of CD137, CD40L or IFN-γ in CTL and Th compartments. (**C–D**) Polyfunctional Th (C) and CTL (D) subset composition represented by pie charts, with overlapping arcs indicating marker co-expression patterns. Significance determined by Friedman’s test; *p < 0·001.

Expansion of vaccine-specific Th cells was dose-dependent in all tested donors for both NEC-T4 and NEC-T5 constructs (10 μg/mL vs. no construct; P = 0·0017 by Friedman’s test; **Figure 4A**). In the CTL compartment, responses were more variable: a clear dose-dependent effect was observed only in expanded PBMCs from donor 215 following NEC-T4 stimulation (LN215; **Figure 4B**, upper panel). In contrast, NEC-T5 more consistently induced dose-dependent CTL responses across donors (P = 0·0069 by Friedman’s test; **Figure 4B**, lower panel).

Despite the lower magnitude of responses observed in some donors—particularly within the CTL compartment—construct-specific polyfunctional responses were consistently detectable in activated cells, primarily at higher vaccine doses. Th cells exhibited substantially higher fractions of stimulus-specific polyfunctional populations. At the 10 μg/mL dose, the fraction of activated Th cells expressing all four AIMs was 16% (LN208), 10% (LN210), and 38% (LN215) for NEC-T4 and 31% (LN208), 20% (LN210), and 10% (LN215) for NEC-T5 (fraction #1; **Figure 4C** and **Supplementary Table S2**). Similarly, the fraction of CD40L⁺CD137⁺TNF⁺ triple-positive Th cells ranged from 39% to 55% for NEC-T4 and from 23% to 49% for NEC-T5 (fraction #3; **Figure 4C** and **Supplementary Table S2**).

Although CTL responses were more limited in magnitude, polyfunctional CTL populations remained detectable. The fraction of activated CTLs co-expressing CD137, IFN-γ, and TNF ranged from 3% to 6% in donors 208 and 210 (fraction #9; **Figure 4D** and **Supplementary Table S2**) and was markedly higher in donor 215 (68% for NEC-T4 and 53% for NEC-T5; fraction #9; **Figure 4D** and **Supplementary Table S2**). Together, these data demonstrate that the immunogenicity of conserved CoV sequences encoded in NEC-T4 and NEC-T5 constructs can be detected *ex vivo* across donors, particularly within the Th compartment.

### mRNA constructs induce robust antigen-specific T-cell responses *in vivo*

The responses to LNP vaccines above were lower than that observed with peptides, possibly due to suboptimal *ex vivo* presentation of vaccine constructs by APC or clonal loss during expansion. Whereas peptide stimulation directly loads MHC molecules, mRNA–LNP vaccines rely on nanoparticle uptake, intracellular trafficking, and antigen expression, processes that are optimized *in vivo* but incompletely recapitulated in standard culture systems [33]. We therefore next evaluated the immunogenicity of NEC-T4 and NEC-T5 in HLA-A2.1 transgenic CB6F1 mice. These mice express a chimeric MHC class I molecule composed of the HLA-A2.1 α1/α2 domains fused to the murine H2-Kb α3 transmembrane and cytoplasmic regions, enabling presentation of human-relevant CTL epitopes. In addition to testing NEC-T4 and NEC-T5, we included a licensed mRNA vaccine (Comirnaty) and two LNP formulation controls: a GFP-LNP vaccine and a spike-LNP vaccine encoding full-length spike (see **Methods**). Mice received one priming intramuscular injection followed by two boosts. Two weeks after the final immunization, splenocytes were harvested and assessed for construct-specific T-cell responses using a triple-color FluoroSpot assay. Cells were re-challenged with overlapping 15-mer peptides spanning the sequences encoded by each construct. For each construct, stimulation peptides were divided into two pools (OVL-4 pools A/B for NEC-T4 and OVL-5 pools A/B for NEC-T5). Similar to the control vaccines (Comirnaty and full spike-LNP), all constructs induced distinct polyfunctional T-cell responses, with clear detection of IFN-γ⁺IL-2⁺TNF⁺ triple-positive cells following re-challenge with the NEC-T4 and NEC-T5 B pools, ranging from 66 ± 19 to 69 ± 30 SFUs (mean ± SD; **Figure 5A**). IL-2 responses followed a similar pattern **(Figure 5B**, upper panel), with B-pool stimulations generally eliciting higher average responses. This suggests that IL-2 is a key contributor to the observed polyfunctional profile and that these cells may possess enhanced proliferative capacity and long-term survival due to IL-2 secretion. Of note, responses to the B pools of NEC-T4 and NEC-T5 overlapped, consistent with the presence of highly immunogenic sequences shared between the constructs (*n* = 29; **Supplementary Table S3**). IFN-γ–producing cells were more abundant than IL-2–producing cells and were clearly antigen-specific for both NEC-T4 and NEC-T5 (1,395 ± 569 to 2,122 ± 114 SFUs; **Figure 5B**, middle panel). However, in contrast to IL-2, IFN-γ production was restricted to stimulation with the B pools, indicating significant differences not only in magnitude but also in the quality of the immunological response. TNF responses followed a pattern similar to IFN-γ, with strong and consistent reactivity upon re-challenge with the B pools, particularly for the NEC-T5 construct (**Figure 5C**, lower panel). Notably, upon rechallenge, the combined OVL pools elicited cytokine release that matched or, in the case of NEC-T5, exceeded that induced by Comirnaty, while maintaining an equivalent polyfunctional (triple-positive) cytokine response (**Supplementary Figure S5**). Collectively, these findings demonstrate that LNP-formulated mRNA constructs encoding conserved CoV sequences elicit strong, specific and physiologically relevant, T-cell responses, supporting their potential as broadly protective immunogens.

**Figure 5.**
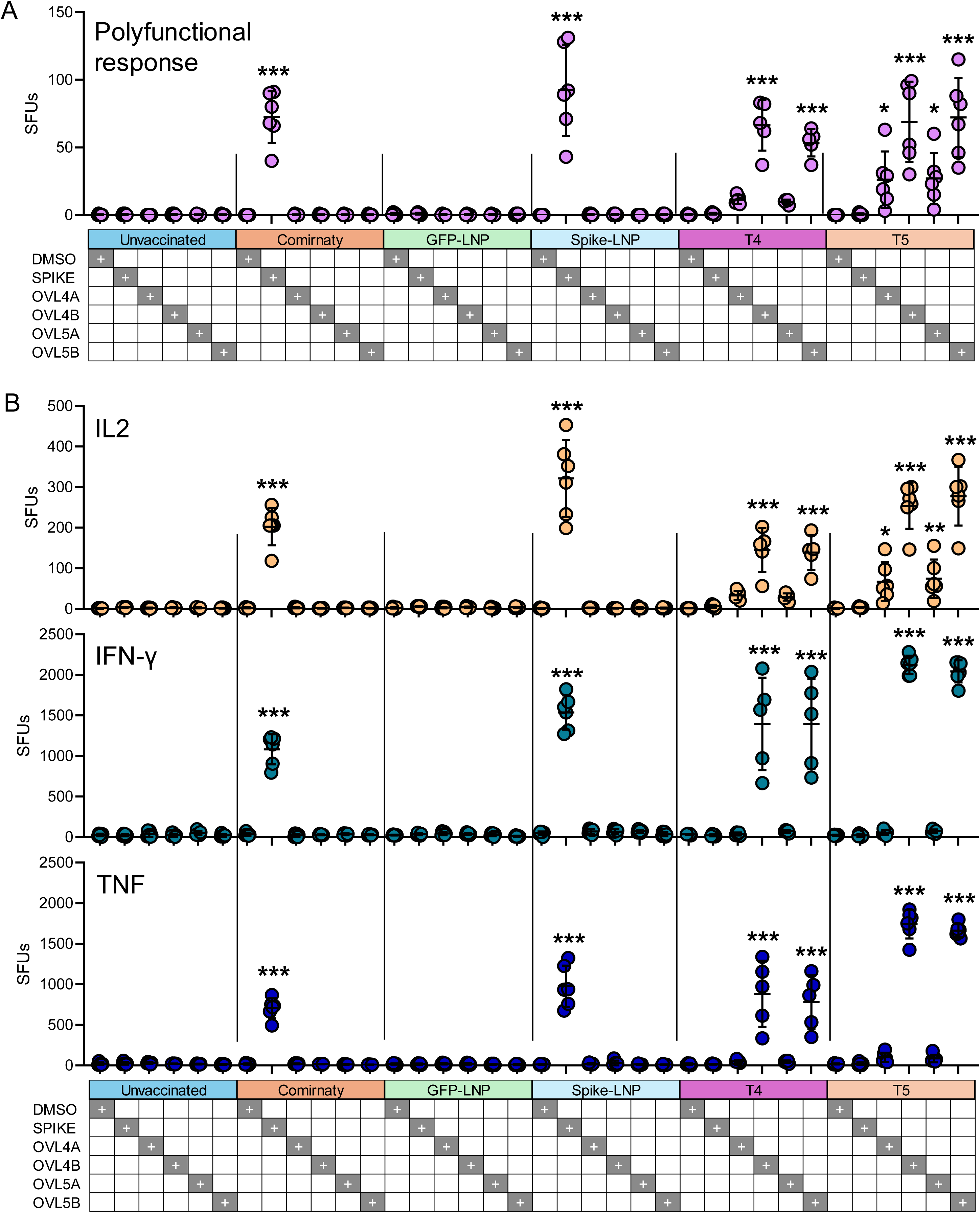
mRNA constructs induce robust antigen-specific T-cell responses *in vivo*. (**A–D**) Reactivity of splenocytes from HLA-A2.1 transgenic CB6F1 mice vaccinated with NEC-T4, NEC-T5, Comirnaty, GFP-LNP, or spike-LNP. Splenocytes were restimulated with overlapping peptides and analyzed by triple-color FluoroSpot. Shown are spot-forming units (SFUs) for (A) triple-positive IFN-γ⁺IL-2⁺TNF⁺, (B) IL-2⁺, IFN-γ⁺, and TNF⁺ T cells. *n* = 6 mice per group; each dot represents the average of two technical replicates. Significance determined by one-way ANOVA with Dunnett’s multiple comparisons test: ***p < 0·0001; **p < 0·02; *p < 0·05.

## Discussion

In this study, we combined machine learning–guided antigen prioritization of conserved T-cell epitopes predicted across the β-CoV proteome with functional immunological validation to identify T-cell targets and evaluate their suitability for multiepitope mRNA vaccination. Antigen selection was performed across all major β-CoV subgenera, encompassing sarbecoviruses as well as embecoviruses, merbecoviruses, nobecoviruses, hibecoviruses, and unclassified β-CoVs, thereby explicitly targeting taxon-level conservation beyond SARS-CoV-2 and its variants.

Across complementary experimental systems, we show that peptide sequences conserved across a greater number of β-CoV taxa were significantly more likely to elicit functional Th and CTL T-cell responses in humans. Conserved peptides induced higher reactivity scores and were more frequently recognized across donors than less conserved counterparts, with a notable enrichment of immunogenic epitopes in non-structural regions. Multiepitope mRNA constructs encoding these conserved regions were efficiently expressed, processed, and presented on HLA class I molecules, as confirmed by immunopeptidomics, and elicited polyfunctional T-cell responses *ex vivo*. *In vivo* vaccination of HLA-A2.1 transgenic mice with these constructs induced robust antigen-specific and polyfunctional T-cell responses comparable in magnitude and quality to those induced by licensed or full-length spike-based mRNA vaccines. Together, these findings establish a direct link between evolutionary conservation and functional T-cell immunogenicity and demonstrate the feasibility of delivering conserved CoV T-cell antigens using an mRNA platform.

The rationale for pursuing conserved T-cell antigens as β-CoV vaccine targets lies in the inherent limitations of spike-centered CoV vaccine strategies for long-term pandemic preparedness. Although current spike-based mRNA vaccines have been highly effective at reducing severe disease, their reliance on neutralizing antibodies directed against a rapidly evolving surface antigen necessitates frequent updating and limits cross-lineage coverage.

In contrast, T-cell immunity targets infected cells through recognition of processed viral peptides, many of which derive from internal or non-structural proteins that are subject to stronger functional constraint and slower evolutionary drift. Importantly, targeting conservation across viral taxa, rather than across variants of a single species, addresses the risk of future zoonotic spillover and enables vaccine designs with relevance beyond the currently circulating human CoVs.

Multiple studies have demonstrated that cross-reactive T-cell immunity toward conserved non-spike epitopes correlates with milder COVID-19 disease or disease avoidance, enhanced viral control in settings of low antibody levels, and durable immune protection [7, 21, 34–38]. A recent extensive study demonstrated that T cells specific for conserved epitopes exhibit significantly enhanced cross-reactivity across multiple β-CoVs, especially for non-spike sequences [39]. Importantly, T cells are necessary for rapid and efficient resolution of COVID-19 [40, 41], for viral control in settings of low antibody levels [42, 43], and to provide durable immunity [44]. Notably, cross-reactivity across β-CoVs in non-spike regions also increases human leukocyte antigen (HLA) coverage relative to spike-focused targets [39]. These considerations support the development of T-cell–focused CoV vaccines to establish a complementary and broadly relevant layer of durable immunity. We found that evolutionary conservation across diverse CoV taxa was a correlate of human T-cell immunogenicity. This finding aligns with previous findings of durable cross reactive T-cell immunity [7, 21, 34–38], an immunity that is expected to accumulate as the population is exposed and re-exposed to β-CoV variants.

Previous vaccine strategies have explored conserved SARS-CoV-2 antigens across variants of concern (VOC), aiming to generate cross-VOC T-cell immunity. This includes 1) an mRNA-LNP construct (BNT162b4) encoding N, M, and short ORF1ab, allowing HLA-A2 bound peptide presentation in mice and protection in hamsters [45], 2) an adenoviral vector (ChAdOx1) delivery of replicase polyproteins (ORF1a/ORF1ab) inducing broad T-cell responses in mice and attenuating disease in hamsters, especially when combined with a spike vaccine [46], 3) a plasmid DNA vaccine that encodes a synthetic chimeric multiepitope protein (SARS-CoV-2 spike, membrane, nucleocapsid, and envelope structural proteins) combining VOC-conserved T-cell regions with overlapping conserved B-cell regions from structural proteins [47], and 4) peptide vaccines with VOC-conserved epitope sets from S2/N/M/E/ORF1ab and adjuvants to drive Th1-biased responses and multi-epitope recognition [48]. While these approaches demonstrate the feasibility of targeting conserved regions within VOC, antigen selection has typically relied on predefined protein choices, sequence conservation alone, or predicted HLA binding affinity.

The present study differs from prior efforts in several important respects. First, antigen prioritization was guided by an integrated machine-learning framework that models antigen processing, HLA presentation, and immunogenicity, and was applied across the entire β-CoV proteome rather than relying solely on sequence conservation or binding affinity. Second, immunogenicity testing identified broadly conserved antigenic sequences mapping to non-structural regions through ex vivo screening of predicted peptides, either individually or in pools. Third, intracellular expression, processing, and HLA class I presentation of encoded antigens were directly validated by global proteomics and immunopeptidomics, thereby bridging computational prediction and biological presentation.

The multiepitope mRNA constructs derived from this prioritization strategy were robustly expressed and processed in vitro and induced polyfunctional T-cell responses in both human *ex vivo* systems and HLA-A2.1 transgenic mice. Although response magnitudes differed between peptide-based stimulation and mRNA delivery in vitro—likely reflecting differences in antigen uptake and processing—the *in vivo* vaccination experiments demonstrated strong, antigen-specific and polyfunctional T-cell responses enriched in IFN-γ, IL-2, and TNF production. These antigen-specific cytokine responses were comparable to, or in the case of NEC-T5 superior to, those induced by licensed spike-based mRNA vaccines, underscoring the potential of conserved T-cell–focused constructs to complement existing vaccine strategies by broadening cellular immune coverage.

This study has several limitations. Although the *ex vivo* vaccination platform allowed functional interrogation of a large number of predicted epitopes across multiple donors, it cannot fully recapitulate the complexity of antigen uptake, tissue distribution, and immune priming that occurs *in vivo*. *In vivo* validation was therefore restricted to HLA-A2.1 transgenic mice, enabling assessment of human-relevant class I–restricted responses but not capturing the full diversity of human HLA backgrounds. In addition, our analyses focused on acute cellular immunogenicity and polyfunctionality rather than durability of memory or protective efficacy following viral challenge. Finally, although conserved epitopes were broadly immunogenic across donors, the cohort size limits conclusions regarding population-level epitope dominance and HLA coverage. These constraints define important directions for future work, including durability studies, heterologous challenge models, and evaluation across expanded HLA repertoires.

In summary, this study demonstrates that evolutionary conservation can be operationalized as a robust and experimentally validated principle for T-cell vaccine antigen selection. By integrating machine-learning–guided prioritization with functional human immune screening, immunopeptidomics, and *in vivo* mRNA vaccination, we provide a scalable framework for identifying and validating conserved viral T-cell targets. Beyond the specific constructs evaluated here, this approach offers a generalizable strategy for the rational design of variant-agnostic T-cell vaccines, with potential applicability to other rapidly evolving viral families and future pandemic threats.

## Supporting information

Supplementary figures description

## Acknowledgements

The authors wish to acknowledge Dr. Javier Castillo-Olivares and Dr. Nadia Cohen of the Coalition for Epidemic Preparedness Innovations (CEPI) for their esteemed guidance and insightful discussions that greatly contributed to the progress and success of this study. We also thank Eline Verheij, Bregtje Smid, and José Harders-Westerveen for the excellent technical execution and appreciate the hard work and dedication of animal biotechnicians and pathology colleagues at the Wageningen Bioveterinary Research. We gratefully acknowledge CEPI for their generous financial support, which made this study possible. Competing interest statement: AO, RiS, SK, BM, MG, PM, RaS, YT, TC and KB were employed by NEC Corporation or its affiliates (NEC Bio, NEC Laboratories Europe) at the time of their contribution to the study.

## References

1. Ruiz-Aravena, M., et al., Ecology, evolution and spillover of coronaviruses from bats. Nature Reviews Microbiology, 2022. 20(5): p. 299–314.

2. Cui, J., F. Li, and Z.-L. Shi, Origin and evolution of pathogenic coronaviruses. Nature Reviews Microbiology, 2019. 17(3): p. 181–192.

3. V’kovski, P., et al., Coronavirus biology and replication: implications for SARS-CoV-2. Nature Reviews Microbiology, 2021. 19(3): p. 155–170.

4. The species Severe acute respiratory syndrome-related coronavirus: classifying 2019-nCoV and naming it SARS-CoV-2. Nat Microbiol, 2020. 5(4): p. 536–544.

5. Ghai, R.R., et al., Animal Reservoirs and Hosts for Emerging Alphacoronaviruses and Betacoronaviruses. Emerg Infect Dis, 2021. 27(4): p. 1015–1022.

6. Pollard, A.J. and E.M. Bijker, A guide to vaccinology: from basic principles to new developments. Nature Reviews Immunology, 2021. 21(2): p. 83–100.

7. Cimen Bozkus, C., et al., T cell epitope mapping reveals immunodominance of evolutionarily conserved regions within SARS-CoV-2 proteome. iScience, 2025. 28(8): p. 113044.

8. Sagar, M., et al., Recent endemic coronavirus infection is associated with less-severe COVID-19. The Journal of Clinical Investigation, 2021. 131(1).

9. Bean, D.J., et al., Heterotypic immunity from prior SARS-CoV-2 infection but not COVID-19 vaccination associates with lower endemic coronavirus incidence. Sci Transl Med, 2024. 16(751): p. eado7588.

10. Tan, C.W., et al., Pan-Sarbecovirus Neutralizing Antibodies in BNT162b2-Immunized SARS-CoV-1 Survivors. N Engl J Med, 2021. 385(15): p. 1401–1406.

11. Dangi, T., et al., Cross-protective immunity following coronavirus vaccination and coronavirus infection. J Clin Invest, 2021. 131(24).

12. Diniz, M.O., et al., Airway-resident T cells from unexposed individuals cross-recognize SARS-CoV-2. Nat Immunol, 2022. 23(9): p. 1324–1329.

13. Loyal, L., et al., Cross-reactive CD4(+) T cells enhance SARS-CoV-2 immune responses upon infection and vaccination. Science, 2021. 374(6564): p. eabh1823.

14. Song, G., et al., Cross-reactive serum and memory B-cell responses to spike protein in SARS-CoV-2 and endemic coronavirus infection. Nature Communications, 2021. 12(1): p. 2938.

15. Lee, R.S.H., et al., Cross-Reactive Antibody Responses to Coronaviruses Elicited by SARS-CoV-2 Infection or Vaccination. Influenza Other Respir Viruses, 2024. 18(5): p. e13309.

16. Kundu, R., et al., Cross-reactive memory T cells associate with protection against SARS-CoV-2 infection in COVID-19 contacts. Nature Communications, 2022. 13(1): p. 80.

17. Lucas, C., et al., Impact of circulating SARS-CoV-2 variants on mRNA vaccine-induced immunity. Nature, 2021. 600(7889): p. 523–529.

18. Krüttgen, A., et al., Large inter-individual variability of cellular and humoral immunological responses to mRNA-1273 (Moderna) vaccination against SARS-CoV-2 in health care workers. Clin Exp Vaccine Res, 2022. 11(1): p. 96–103.

19. Narowski, T.M., et al., SARS-CoV-2 mRNA vaccine induces robust specific and cross-reactive IgG and unequal neutralizing antibodies in naive and previously infected people. Cell Reports, 2022. 38(5).

20. Liu, M.Q., et al., Inactivated SARS-CoV-2 Vaccine Shows Cross-Protection against Bat SARS-Related Coronaviruses in Human ACE2 Transgenic Mice. J Virol, 2022. 96(8): p. e0016922.

21. Mateus, J., et al., Selective and cross-reactive SARS-CoV-2 T cell epitopes in unexposed humans. Science, 2020. 370(6512): p. 89–94.

22. Swadling, L., et al., Pre-existing polymerase-specific T cells expand in abortive seronegative SARS-CoV-2. Nature, 2022. 601(7891): p. 110–117.

23. Amoutzias, G.D., et al., The Remarkable Evolutionary Plasticity of Coronaviruses by Mutation and Recombination: Insights for the COVID-19 Pandemic and the Future Evolutionary Paths of SARS-CoV-2. Viruses, 2022. 14(1).

24. Kandwal, S. and D. Fayne, Genetic conservation across SARS-CoV-2 non-structural proteins - Insights into possible targets for treatment of future viral outbreaks. Virology, 2023. 581: p. 97–115.

25. Federico, L., et al., Experimental validation of immunogenic SARS-CoV-2 T cell epitopes identified by artificial intelligence. Frontiers in Immunology, 2023. **Volume** 14 - 2023.

26. Malone, B., et al., Artificial intelligence predicts the immunogenic landscape of SARS-CoV-2 leading to universal blueprints for vaccine designs. Sci Rep, 2020. 10(1): p. 22375.

27. De Keersmaecker, B., et al., Lumenal part of the DC-LAMP protein is not required for induction of antigen-specific T cell responses by means of antigen-DC-LAMP messenger RNA-electroporated dendritic cells. Hum Gene Ther, 2010. 21(4): p. 479–85.

28. Federico, L., et al., Robust spike-specific CD4+ and CD8+ T cell responses in SARS-CoV-2 vaccinated hematopoietic cell transplantation recipients: a prospective, cohort study. Frontiers in Immunology, 2023. **Volume** 14 - 2023.

29. Roederer, M., J.L. Nozzi, and M.C. Nason, SPICE: Exploration and analysis of post-cytometric complex multivariate datasets. Cytometry Part A, 2011. 79A(2): p. 167–174.

30. ; Available from: https://niaid.github.io/spice.

31. Gainullin, M., et al., People who use drugs show no increase in pre-existing T-cell cross-reactivity toward SARS-CoV-2 but develop a normal polyfunctional T-cell response after standard mRNA vaccination. Front Immunol, 2023. 14: p. 1235210.

32. Gordon, J.L., et al., Development of broadly protective coronavirus vaccines: A joint NIAID-CEPI workshop report. Vaccine, 2025. 54: p. 126909.

33. Voke, E., et al., Protein corona formed on lipid nanoparticles compromises delivery efficiency of mRNA cargo. Nat Commun, 2025. 16(1): p. 8699.

34. Mallajosyula, V., et al., CD8+ T cells specific for conserved coronavirus epitopes correlate with milder disease in patients with COVID-19. Science Immunology, 2021. 6.

35. Kundu, R., et al., Cross-reactive memory T cells associate with protection against SARS-CoV-2 infection in COVID-19 contacts. Nature Communications, 2022. 13.

36. Meyer, S., et al., Prevalent and immunodominant CD8 T cell epitopes are conserved in SARS-CoV-2 variants. Cell Rep, 2023. 42(1): p. 111995.

37. Saini, S.K., et al., SARS-CoV-2 genome-wide T cell epitope mapping reveals immunodominance and substantial CD8(+) T cell activation in COVID-19 patients. Sci Immunol, 2021. 6(58).

38. Lineburg, K.E., et al., CD8(+) T cells specific for an immunodominant SARS-CoV-2 nucleocapsid epitope cross-react with selective seasonal coronaviruses. Immunity, 2021. 54(5): p. 1055–1065.e5.

39. Pereira Neto, T.A., et al., Highly conserved Betacoronavirus sequences are broadly recognized by human T cells. Cell, 2025. 188(20): p. 5653–5665.e12.

40. Rydyznski Moderbacher, C., et al., Antigen-Specific Adaptive Immunity to SARS-CoV-2 in Acute COVID-19 and Associations with Age and Disease Severity. Cell, 2020. 183(4): p. 996–1012.e19.

41. Tan, A.T., et al., Early induction of functional SARS-CoV-2-specific T cells associates with rapid viral clearance and mild disease in COVID-19 patients. Cell Rep, 2021. 34(6): p. 108728.

42. McMahan, K., et al., Correlates of protection against SARS-CoV-2 in rhesus macaques. Nature, 2021. 590(7847): p. 630–634.

43. Swadling, L., et al., Pre-existing polymerase-specific T cells expand in abortive seronegative SARS-CoV-2. Nature, 2021.

44. Le Bert, N., et al., SARS-CoV-2-specific T cell immunity in cases of COVID-19 and SARS, and uninfected controls. Nature, 2020. 584(7821): p. 457–462.

45. Christina, M.A., et al., The T-cell-directed vaccine BNT162b4 encoding conserved non-spike antigens protects animals from severe SARS-CoV-2 infection. Cell, 2023. 186: p. 2392 – 2409.e21–2392 – 2409.e21.

46. Wee, E., et al., Design, Immunogenicity and Preclinical Efficacy of the ChAdOx1.COVconsv12 Pan-Sarbecovirus T-Cell Vaccine. Vaccines, 2024. 12.

47. Lauren, M.M., et al., Highly conserved, non-human-like, and cross-reactive SARS-CoV-2 T cell epitopes for COVID-19 vaccine design and validation. NPJ Vaccines, 2021. 6.

48. Prakash, S., et al., Cross-protection induced by highly conserved human B, CD4+, and CD8+ T-cell epitopes-based vaccine against severe infection, disease, and death caused by multiple SARS-CoV-2 variants of concern. Frontiers in Immunology, 2024. 15.

