## Supplementary figures description for "Machine Learning–Driven Antigen Selection Reveals Conserved T-Cell Targets for Broad Coronavirus Vaccination"

**Supplementary Table S1. Complete list of peptides included in the pools and RNA constructs used in the study**. For each peptide, the table lists the sequence identifier (Seq ID No.), sequence length, protein region, pool assignment (B1–B3), construct assignment (NEC-T4–T6), testing context, detectability by mass spectrometry in immunopeptidomic analyses, and the number and identity of covered subgenera and species as well as immunogenicity and binding scores across selected HLA alleles. Peptides whose reactivity scores were used to generate the heatmaps of Supplementary Figure S2 are identified in column G. Peptide immunogenicity and binding scores were generated using the NEC Immune Profiler as described in Methods.

**Supplementary Figure S1. Identification of conserved epitopes for T- cell response analyses.** (**A**). Individual peptides analyzed in the study (listed in Supplementary Table S1) plotted as a function of the number of β-CoV subgenera (sarbecovirus, embecovirus, merbecovirus, nobecovirus, hibecovirus, and unclassified β-CoVs ) and the number of distinct coronavirus taxa (defined as individual viral isolates or reference sequences) in which they are found. (**B**) β-CoV subgenera–specific conservation. Bars indicate the number of peptides present in each combination of subgenera. (**C**) Sarbecovirus-specific conservation. Bars indicate the number of peptides present in each combination of SARS-CoV (SARS-CoV-1 ∪ SARS-CoV-2), bat, and pangolin CoVs.

**Supplementary Figure S2. T cells display greater likelihood of reactivity toward shared epitopes.** Heatmap of reactivity scores for individually tested peptides from 7 healthy donors in both CTL and Th compartments. The degree of sharing (number of species/subgenera in which the epitope appears) is indicated. Peptides are sorted in descending order of species coverage (236 to 1). The horizontal line marks the cut-off for shared epitopes (5 species).

**Supplementary Table S2. Frequencies of polyfunctional T-cell populations following *ex vivo* vaccination.** Comprehensive list of polyfunctional subsets and their relative abundance after stimulation with B1, B2, B3, NEC-T4, and NEC-T5 across CTL and Th compartments.

**Supplementary Figure S3. Response to *ex vivo* vaccination with SARS-CoV-2 peptides is detectable and specific** (**A**) Schematic of the PBMC *ex vivo* vaccination workflow using antigenic peptide pools (B1, B2, and Peptivator). After initial stimulation, T cells were re-challenged twice (days 15 and 25) and analysed by flow cytometry on day 35. (**B**) Representative examples of antigen-specific T-cell expansion following stimulation. Activation was measured as the percentage of T cells expressing CD137, CD40L, and IFN-γ in both CTL and Th compartments. (**C–D**) Quantification of antigen-specific reactive CTL (C) and Th (D) cells from 3 donors at day 35. (**E–F**) Summary of stimulus-specific responses across a cohort of 17 healthy donors for CTL (E) and Th (F). Data are presented as in C–D after background subtraction.

**Supplementary Table S3. List of NEC-T4 and NEC-T5 epitope hotspots and corresponding peptides used for *in vivo* re-challenge pools.**

**Supplementary Figure S4. Mapping of expressed pMHC-I peptides and accuracy of immunogenicity predictions.** (**A**) Peptides identified by mass spectrometry from MHC-I complexes isolated 48 h post-transfection. Peptides are mapped to construct position (x-axis) with predicted immunogenicity scores (y-axis), color-coded by HLA allele. (**B**) β-CoV subgenus coverage by identified peptides. Bars represent the proportion of species covered out of the total predicted; absolute numbers are shown above each bar. (**C–D**) ROC curves showing prediction accuracy of immunogenicity scores for (C) all HLAs combined and (D) individual HLAs (HLA-A02:01, HLA-A03:01, HLA-B*07:02). AUC values indicated.

**Supplementary Figure S5. Antigen-specific cytokine responses to mRNA constructs and the Pfizer–BioNTech COVID-19 vaccine.** Spot-forming units (SFUs) are shown for IFN-γ⁺, TNF⁺, IL-2⁺, and triple-positive IFN-γ⁺IL-2⁺TNF⁺ (Triple⁺) T cells. *n* = 6 mice per group; each dot represents the mean of two technical replicates. Statistical significance was determined by one-way ANOVA with Dunnett’s multiple comparisons test; p values are indicated in the figure.
