## Supplemental Figures for "Machine Learning–Driven Antigen Selection Reveals Conserved T-Cell Targets for Broad Coronavirus Vaccination"

**Supplementary Figure S1.** Identification of conserved epitopes for T- cell response analyses.

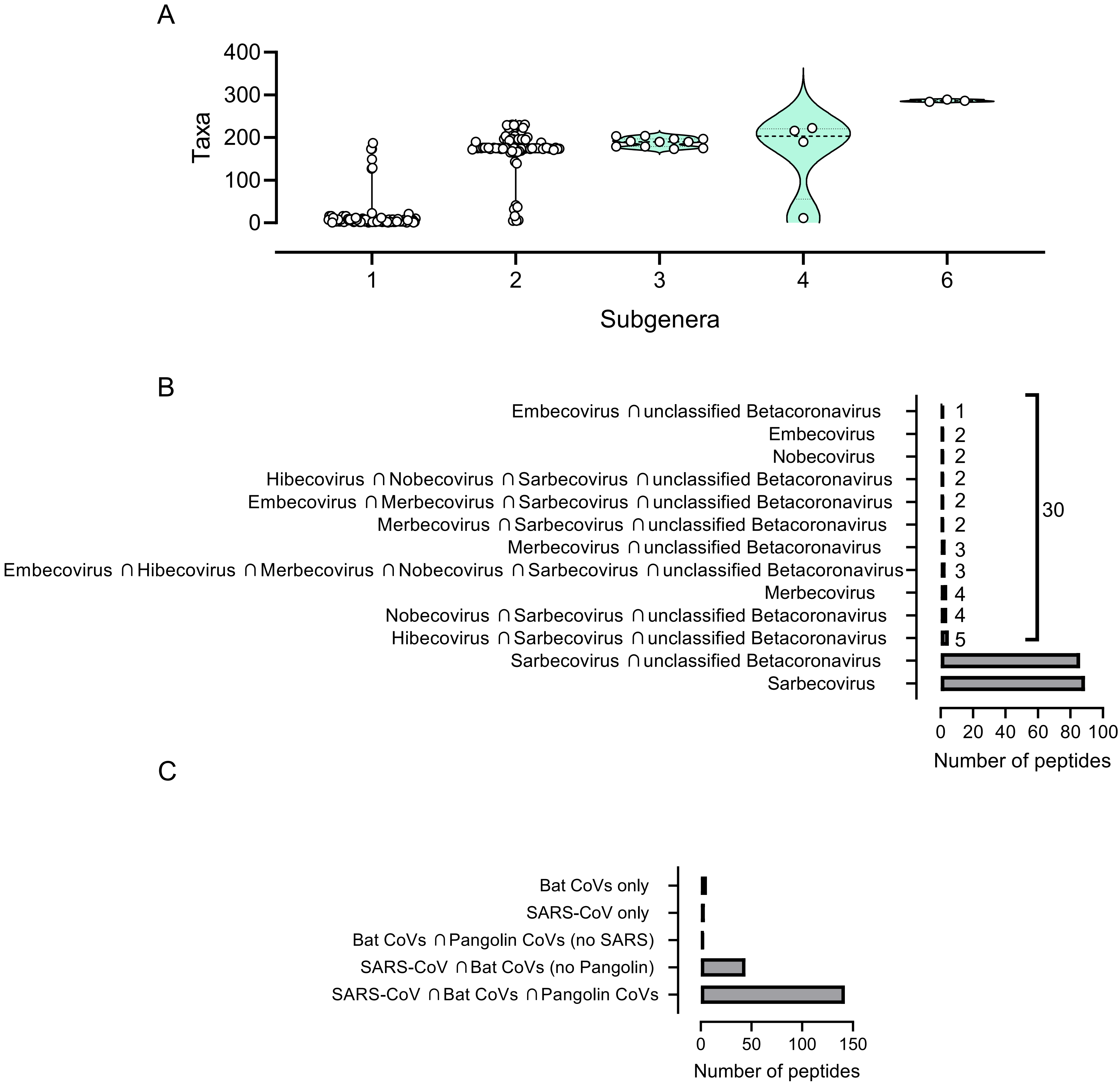

**Supplementary Figure S2.** T cells display greater likelihood of reactivity toward shared epitopes.

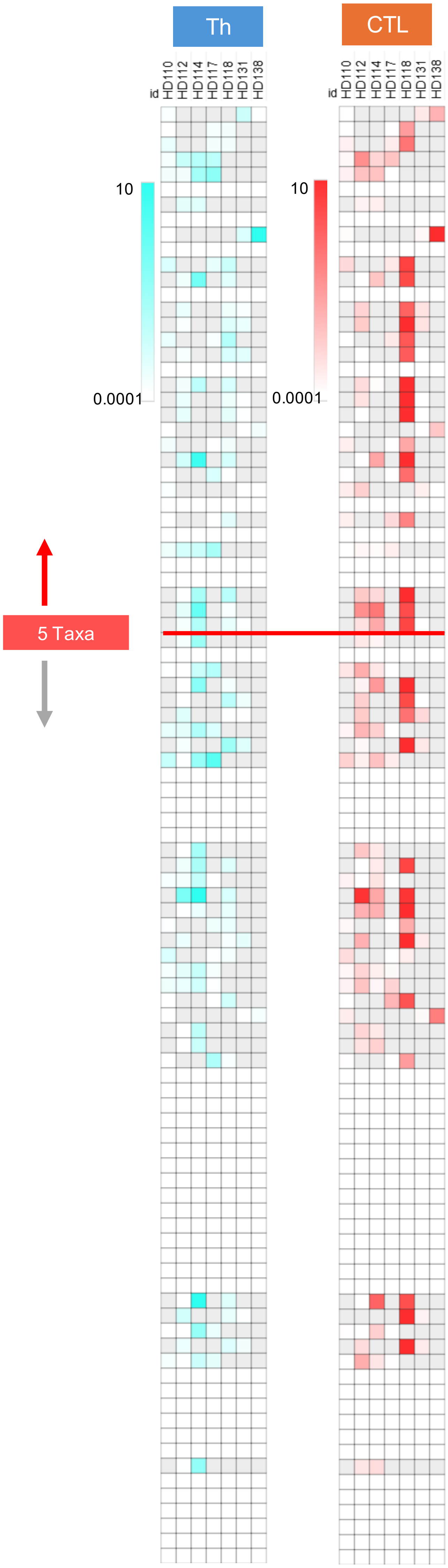


| No. Taxa | No. Subgen | Peptide Seq. | Region |
|---|---|---|---|
| 236 | 6 | RARTVAGVSI | Orf1ab |
| 235 | 6 | MNLKYAISA | Orf1ab |
| 234 | 6 | TQMNLKYAI | Orf1ab |
| 177 | 2 | ILTALRLCAY | E |
| 176 | 2 | MAYRFNGIG | S |
| 176 | 2 | RLITGRLQSL | S |
| 175 | 2 | FAMQMAYRF | S |
| 174 | 2 | IPFAMQMAY | S |
| 172 | 2 | RAAEIRASA | S |
| 172 | 2 | AEIRASANL | S |
| 169 | 2 | VVRQCSGVTF | Orf1ab |
| 169 | 2 | ALYNKYKYF | Orf1ab |
| 169 | 2 | FLLNKEMYL | Orf1ab |
| 168 | 2 | YMPASWVMRI | Orf1ab |
| 168 | 2 | MPASWVMRIM | Orf1ab |
| 168 | 2 | TQYNRYLALY | Orf1ab |
| 168 | 2 | TQYNRYLAL | Orf1ab |
| 168 | 2 | MPASWVMRI | Orf1ab |
| 167 | 2 | HFISNSWLM | Orf1ab |
| 167 | 2 | SWVMRIMTW | Orf1ab |
| 167 | 2 | KFYGGWHNM | Orf1ab |
| 167 | 2 | YEDQDALFAY | Orf1ab |
| 167 | 2 | LLPSLATVAY | Orf1ab |
| 166 | 2 | MSYEDQDALF | Orf1ab |
| 165 | 1 | NMVYMPASW | Orf1ab |
| 165 | 1 | TVAYFNMVY | Orf1ab |
| 163 | 2 | FAYTKRNVI | Orf1ab |
| 158 | 2 | LNKEMYLKL | Orf1ab |
| 135 | 1 | YQPYRVVVL | S |
| 39 | 2 | FPREGVFVSN | S |
| 30 | 2 | KTSVDCTMY | S |
| 10 | 6 | IKWPWYIWL | S |
| 6 | 1 | APISAMVRMY | Orf1ab |
| 6 | 1 | APISAMVRM | Orf1ab |
| 6 | 1 | VQMAPISAM | Orf1ab |
| 5 | 1 | RGWIFGTTL | S |
| 5 | 1 | AHFPREGVF | S |
| 4 | 1 | IIMRLWLCWK | Orf3a |
| 4 | 1 | AMMFVKHKH | Orf1ab |
| 4 | 1 | VMFTPLVPF | Orf1ab |
| 4 | 1 | MMFVKHKHA | Orf1ab |
| 4 | 1 | FLYLYALVYF | Orf3a |
| 4 | 1 | WLIINLVQM | Orf1ab |
| 4 | 1 | GLAAIMQLFF | Orf1ab |
| 4 | 1 | SLVKPSFYVY | E |
| 4 | 1 | MVMFTPLVPF | Orf1ab |
| 4 | 1 | LLYDANYFL | Orf3a |
| 4 | 1 | WLMWLIINL | Orf1ab |
| 4 | 1 | MFVKHKHAF | Orf1ab |
| 3 | 1 | VYYPDKVFRS | S |
| 3 | 1 | EMIAQYTSAL | S |
| 3 | 1 | SNIIRGWIF | S |
| 3 | 1 | HFYWFFSNY | Orf1ab |
| 3 | 1 | AMSAFAMMF | Orf1ab |
| 3 | 1 | STKHFYWFF | Orf1ab |
| 3 | 1 | IAMSAFAMM | Orf1ab |
| 3 | 1 | LFFSYFAVHF | Orf1ab |
| 3 | 1 | KHFYWFFSNY | Orf1ab |
| 3 | 1 | AIMQLFFSYF | Orf1ab |
| 3 | 1 | LEAPFLYLYA | Orf3a |
| 3 | 1 | SINFVRIIM | Orf3a |
| 3 | 1 | FHLDGEVITF | Orf1ab |
| 3 | 1 | RTIKGTHHW | Orf1ab |
| 3 | 1 | REVRTIKVF | Orf1ab |
| 3 | 1 | GEVITFDNL | Orf1ab |
| 3 | 1 | LEAPFLYLY | Orf3a |
| 3 | 1 | AIMQLFFSY | Orf1ab |
| 3 | 1 | YLKLRSDVL | Orf1ab |
| 3 | 1 | GTHWFVTQR | S |
| 3 | 1 | FVSNGTHWFV | S |
| 3 | 1 | APFLYLYAL | Orf3a |
| 3 | 1 | TRFQTLLAL | S |
| 3 | 1 | IPTNFTISV | S |
| 3 | 1 | LPIGINITRF | S |
| 3 | 1 | SIIAYTMSL | S |
| 3 | 1 | TLLALHRSY | S |
| 3 | 1 | FKNLREFVF | S |
| 3 | 1 | GAAAYYVGY | S |
| 3 | 1 | LPFNDGVYF | S |
| 2 | 1 | WTAGAAAYYV | S |
| 2 | 1 | VYYTSNPTTF | Orf1ab |
| 2 | 1 | DMVDTSLSGF | Orf1ab |
| 2 | 1 | MGIIAMSAF | Orf1ab |
| 2 | 1 | FVFKNIDGYF | S |
| 2 | 1 | LPFAMGIIAM | Orf1ab |
| 2 | 1 | TLKSFTVEK | S |
| 2 | 1 | YLQPRTFLL | S |
| 2 | 1 | WTAGAAAYY | S |
| 2 | 1 | DKVFRSSVL | S |
| 2 | 1 | FVFKNIDGY | S |
| 1 | 1 | IAIVMVTIM | S |
| 1 | 1 | TRFASVYAW | S |
| 1 | 1 | ESIVRFPNI | S |
| 1 | 1 | NSFTRGVYY | S |
| 1 | 1 | YTNSFTRGVY | S |
| 1 | 1 | KVGGNYNYLY | S |
| 1 | 1 | TPINLVRDL | S |

**Supplementary Figure S3.** Response to ex vivo vaccination with β-CoV peptides is detectable and specific.

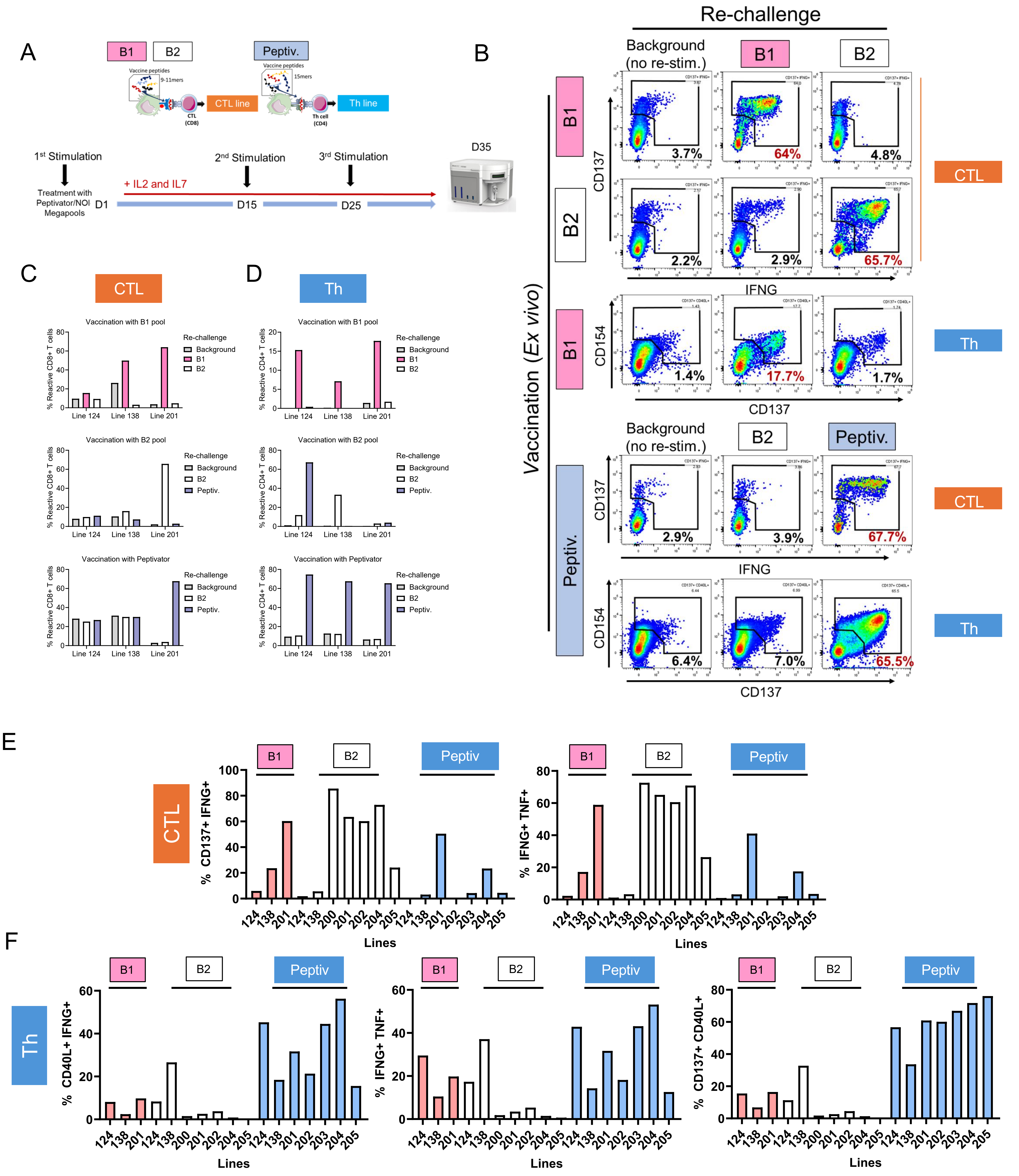

**Supplementary Figure S4.** Mapping of expressed pMHC-I peptides and accuracy of immunogenicity predictions.

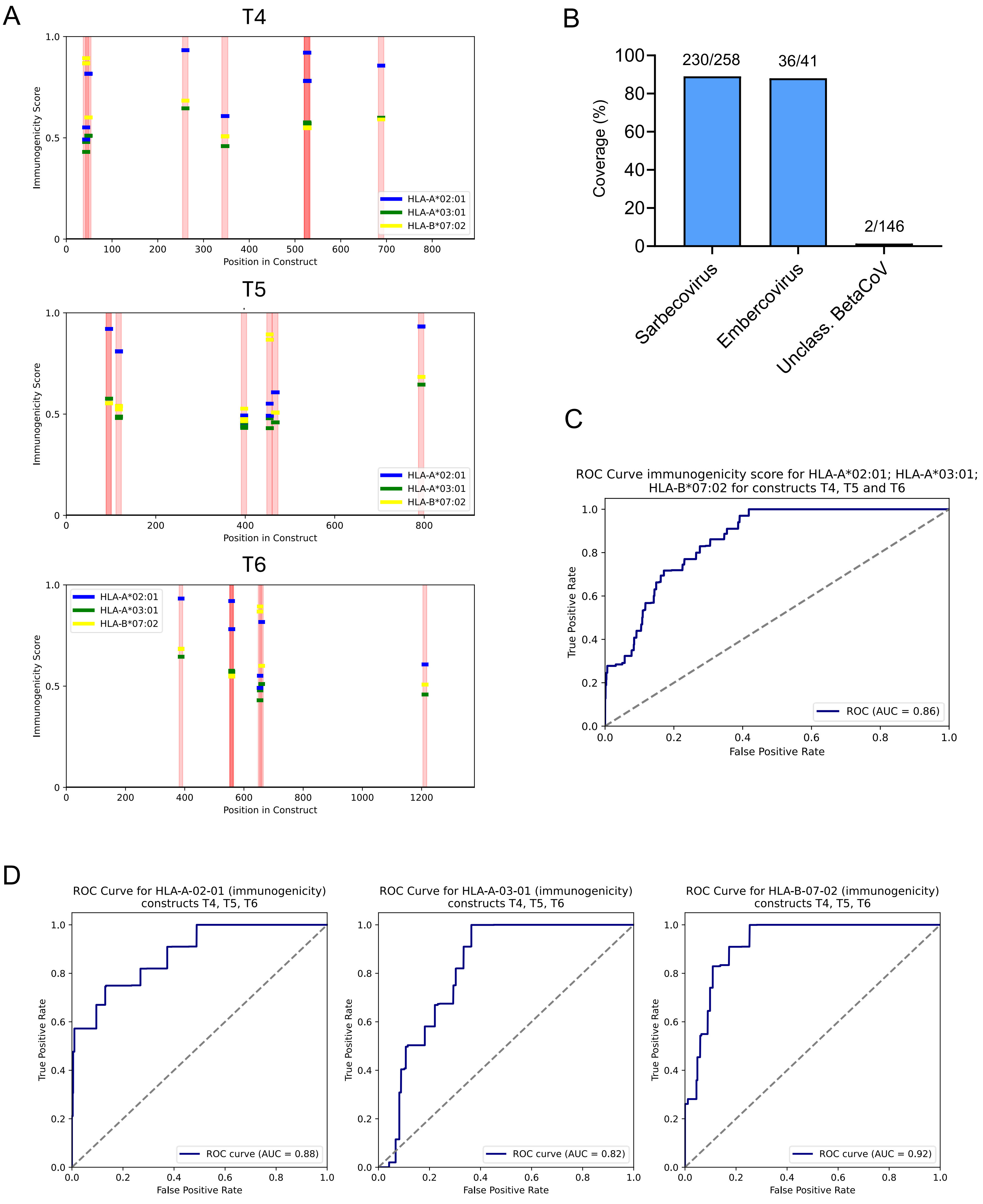

**Supplementary Figure S5.** Antigen-specific cytokine responses to mRNA constructs and the Pfizer–BioNTech COVID-19 vaccine.

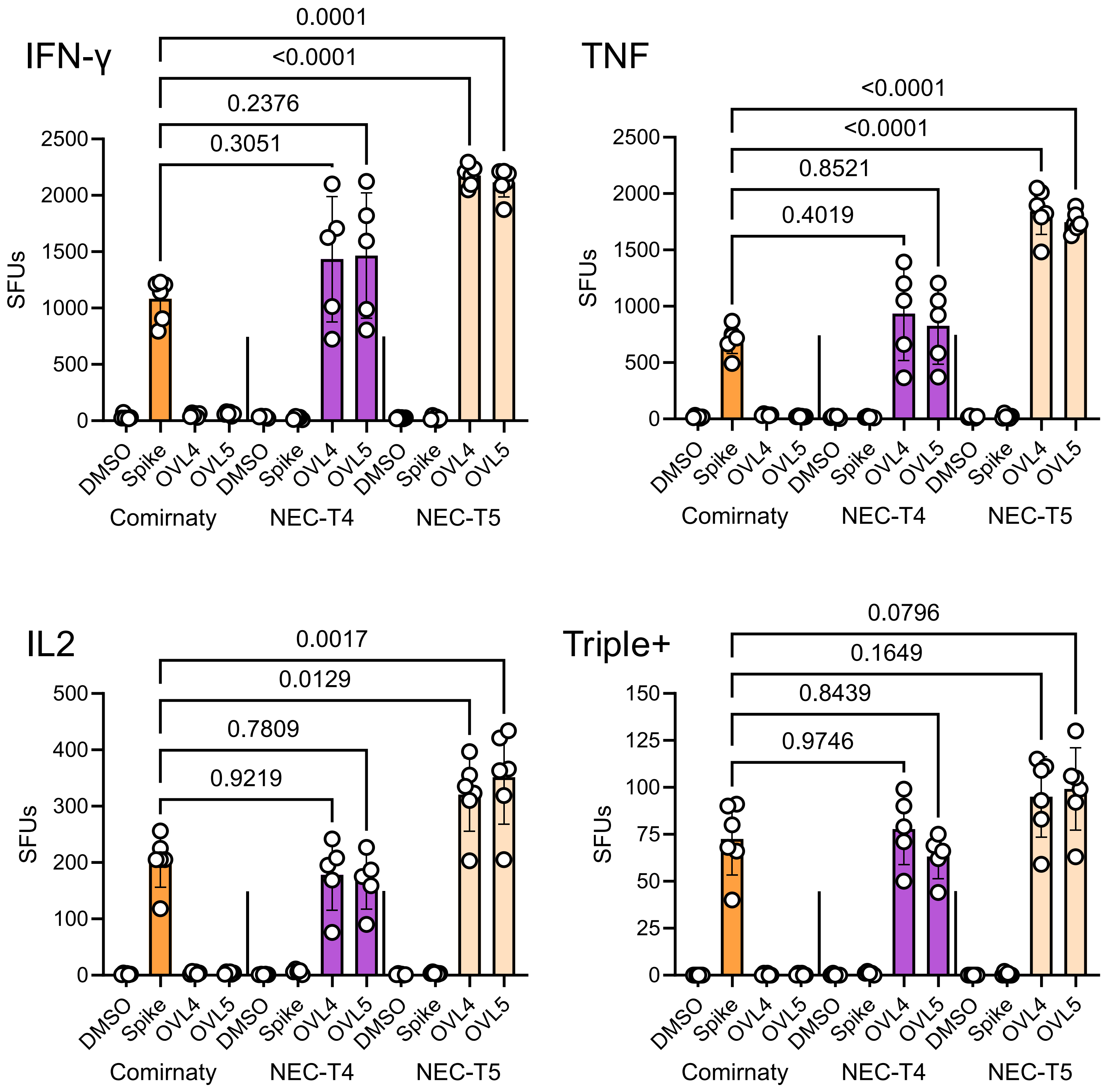
